# Proteolytic Remodeling of Cargo Receptor Networks by RHBDL4 Tunes Secretory Pathway Flux

**DOI:** 10.64898/2026.08.25.746970

**Authors:** Susanne S. Steigleder, Clara Neumann, Marina Tauber, Isabel Krämer, Monika Pesch, Julia D. Knopf, Julian Nüchel, Marius K. Lemberg

## Abstract

Cargo receptors are central organizers of the secretory pathway, yet the mechanisms controlling their abundance remain poorly understood. The endoplasmic reticulum (ER)-resident intramembrane protease RHBDL4 promotes substrate turnover via a non-canonical branch of ER-associated degradation and has recently been implicated in regulating secretory pathway components. We previously identified the p24 cargo receptor TMED7 as an RHBDL4 substrate, suggesting that cargo receptor turnover contributes to secretory pathway regulation. Here, quantitative proteomics identify members of the ER-Golgi intermediate compartment (ERGIC) cargo receptor family as endogenous RHBDL4 substrates, demonstrating that RHBDL4 targets multiple cargo receptor families within the early secretory pathway. Accordingly, RHBDL4 modulates multiple ERGIC-dependent transport pathways. In addition, unbiased secretome analysis reveals increased secretion of lysosomal precursor proteins upon RHBDL4 ablation. Mechanistically, we show that this phenotype is mediated, at least in part, by RHBDL4-dependent cleavage of the lysosomal cargo receptor sortilin/SORT1. Together, these findings identify cargo receptors as a major class of RHBDL4 substrates and establish proteolytic remodeling of cargo receptor networks as a mechanism for regulating secretory pathway flux.

## Introduction

The secretory pathway delivers proteins to the endo-lysosomal system, plasma membrane, and extracellular space and therefore requires tight regulation to ensure appropriate protein localization and abundance. Within this pathway vesicular transport connects organelles and enables selective cargo sorting (Bednarek *et al*, 1996). While some proteins move by bulk flow, most rely on coat protein complex II (COPII) components for export from the endoplasmic reticulum (ER) to the Golgi apparatus. Soluble or transmembrane (TM) proteins that lack COPII-interacting motifs require cargo receptors to engage the transport machinery (Barlowe *et al*, 1994; Farhan *et al*, 2025). Several cargo receptor families have evolved to accommodate the remarkable diversity of proteins traversing the secretory pathway, including the p24/TMED proteins, the LMAN cargo receptor family, and the ER-Golgi intermediate compartment (ERGIC) cargo receptors (Adolf *et al*, 2019; Satoh *et al*, 2014; Strating & Martens, 2009). Among these, ERGIC cargo receptors occupy a central position at the interface between the ER and Golgi apparatus (Adolf *et al*., 2019; Yang *et al*, 2026). The ERGIC family comprises three members involved in bidirectional ER-Golgi transport and plays key roles in regulating transport between early secretory compartments, transporting cargo as diverse as Wnt receptor Evi/WLS and haptoglobin (Yoo *et al*, 2019; Yu *et al*, 2014). These receptors share a conserved architecture, consisting of a large luminal cargo-binding domain flanked by two TM helices (Breuza *et al*, 2004; Moreau *et al*, 2011; Orci *et al*, 2003). The coexistence of multiple cargo receptor families raises the question of how cells dynamically adjust cargo receptor abundance to meet changing secretory demands while maintaining trafficking fidelity.

Following transit through the Golgi, cargo proteins are sorted at the trans-Golgi network to their final destinations (Ford *et al*, 2021). Most lysosomal proteins are tagged with mannose-6-phosphate (M6P), which is recognized by M6P receptors that mediate endosomal delivery (Braulke & Bonifacino, 2009). However, M6P-independent trafficking routes also exist, including sortilin/SORT1-mediated delivery of selected lysosomal proteins (Canuel *et al*, 2008). Sortilin binds cargo through its luminal Vps10 domain and recruits clathrin adaptor proteins via cytosolic motifs (reviewed in (Mitok *et al*, 2022)). This pathway is best characterized for prosaposin (PSAP), a precursor of saposins that facilitate glycosphingolipid degradation (Lefrancois *et al*, 2003), but sortilin has also been implicated in the trafficking of selected cathepsins (Coutinho *et al*, 2012). Together, these pathways illustrate the complexity of cargo receptor networks and highlight the need for mechanisms that dynamically remodel cargo receptor abundance in response to changing trafficking demands.

An emerging mechanism for regulating secretion flux involves selective turnover of trafficking regulators and COPII components (Avci *et al*, 2019; Omari *et al*, 2018; Ossareh-Nazari *et al*, 2010; Wunderle *et al*, 2016). Recently, we showed that the intramembrane protease RHBDL4 regulates members of the p24/TMED cargo receptor family (Knopf *et al*, 2024), providing the first evidence that cargo receptor abundance is actively controlled through proteolytic turnover rather than solely by biosynthesis and constitutive degradation. Through this mechanism, RHBDL4 limits TMED7-dependent TLR4 trafficking and suppresses excessive innate immune signaling, thereby establishing cargo receptor quantity control as a regulatory principle for secretion. These findings further extend the previously identified role of RHBDL4 in ER quality control as a non-canonical ER-associated degradation (ERAD) factor that promotes proteolytic turnover via p97/VCP-dependent extraction (Bock *et al*, 2022; Fleig *et al*, 2012). This raises the possibility that RHBDL4 functions as a broader regulator of cargo receptor network composition, rather than a protease dedicated to individual trafficking pathways. Supporting this idea, in this study, we identify ERGIC cargo receptors and the lysosomal sorting receptor sortilin as additional RHBDL4 substrates and demonstrate that RHBDL4 modulates multiple trafficking pathways involving these factors. Our results support a unifying model in which proteolytic tuning of cargo receptor networks enables rapid remodeling of secretory pathway architecture, allowing cells to selectively redirect protein trafficking without globally altering secretion.

## Results

### Identification of ERGIC cargo receptors as RHBDL4 substrates

Using stable-isotope labeling by amino acids in cell culture (SILAC)-based quantitative proteomics in RHBDL4 knockout cells, we previously identified the p24 family proteins TMED3, TMED6, and TMED7 as RHBDL4 substrates, implicating the protease in the regulation of secretion dynamics (Knopf *et al*., 2024). To explore whether this regulatory role extends to other trafficking factors, we performed a second quantitative proteomic screen. Strikingly, two members of ERGIC cargo receptor family, ERGIC1 and ERGIC3, emerged as candidate RHBDL4 substrates (**Fig. 1A** and **Table EV1**). To validate this, we performed rhomboid cleavage assays using the three ERGIC family members (ERGIC1-3), co-expressed as N-terminally FLAG-tagged constructs together with either wild-type RHBDL4 or its catalytically inactive S144A mutant (Fleig *et al*., 2012). Cleavage fragment generation was strictly dependent on RHBDL4 activity (**Figs. 1B** and **EV1A-B**), and quantification revealed ERGIC2 as the most efficiently cleaved ERGIC protein, while ERGIC1 and ERGIC3 showed lower processing efficiency (**Fig. 1C**). Consistent with RHBDL4-dependent degradation via the ERAD pathway, inhibition of either the proteasome or p97 stabilized RHBDL4-generated fragments (**Figs. 1B** and **EV1A-E**).

**Figure 1.**
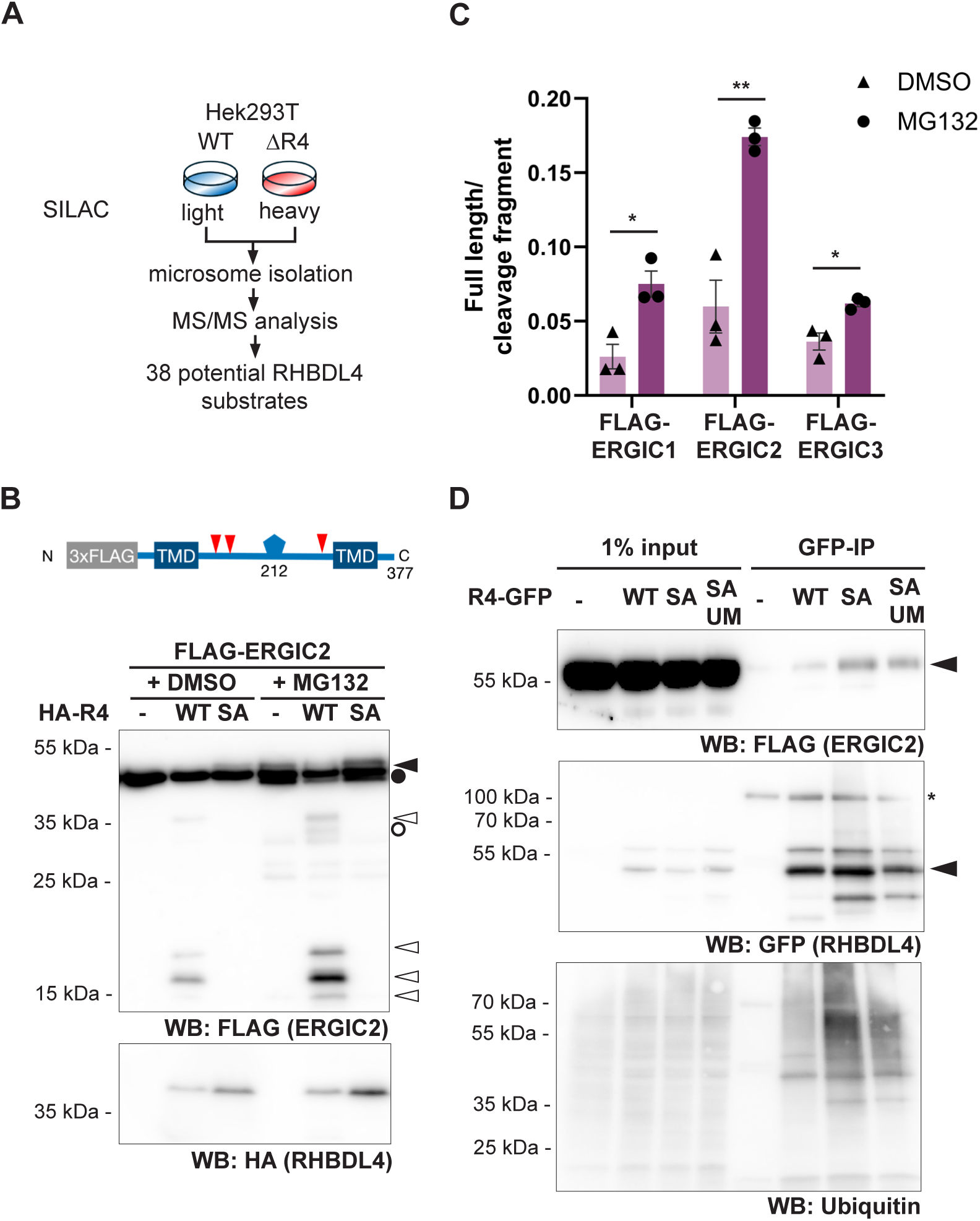
RHBDL4 cleaves members of the ERGIC but not the LMAN cargo receptor family. (**A**)Experimental outline of RHBDL4 substrate screen in HEK293T wild-type (WT) and RHBDL4 knockout (ΔR4) cells. Cells were SILAC-labeled and microsomes were isolated by differential centrifugation. Samples were analyzed by tandem mass spectrometry (MS/MS). (**B**) Western blot (WB) analysis of HEK293T cells expressing FLAG-ERGIC2 (closed arrow) together with either empty vector (-), wild-type (WT) HA-RHBDL4 (HA-R4), or the catalytically inactive SA mutant. Co-expression of HA-R4 WT generates N-terminal cleavage fragments of approximately 35, 20, 18, and 15 kDa (open arrows), which are stabilized by MG132 treatment (2 µM; DMSO, vehicle control). Upper panel, schematic of the FLAG-ERGIC2 construct. TMD, TM domain. **(C)** Quantification of the cleavage assays shown in (B) and Figure EV1A-B based on the ratio of full-length substrate to the summed intensity of the cleavage fragments (means ± SEM, n = 3, *p < 0.05, **p < 0.01; unpaired two-sided Student’s t-test). **(D)** HEK293T cells expressing chromosomally FLAG-tagged ERGIC2 were transiently transfected with either empty vector, WT GFP-tagged RHBDL4 (R4-GFP), the catalytically inactive SA mutant or the SA-UIM double mutant (SAUM), were lysed with Triton X-100 and subjected to GFP-specific immunoprecipitation (IP). Cells were treated with CB-5083 (2.5 µM) prior to lysis. Background bands are marked with an asterisk.

### Endogenous substrate trapping confirms ERGIC2 and ERGIC3 as RHBDL4 substrates

Having established RHBDL4-dependent cleavage of all three ERGIC family members in overexpression assays, we next asked whether endogenous ERGIC proteins engage RHBDL4 under physiological conditions. Given the known difficulty in detecting RHBDL4-generated cleavage fragments at endogenous levels (Knopf *et al*., 2024), we tested for functional interaction with native ERGIC2 and ERGIC3, both of which have defined cargo (Guan *et al*, 2022; Moreau *et al*., 2011; Yang *et al*., 2026; Yoo *et al*., 2019; Yu *et al*., 2014). We previously demonstrated that the catalytically inactive S144A RHBDL4 mutant (R4-SA) can trap substrates through stable interactions, reflecting a stalled proteolytic interaction. This substrate-trapping behavior is reduced in a double mutant additionally lacking a functional ubiquitin-interacting (R4-SAUM), which contributes to substrate recruitment (Fleig *et al*., 2012; Knopf *et al*, 2020).

Consistent with physiological substrate engagement, both endogenously FLAG-tagged ERGIC2 and endogenous, untagged ERGIC3 efficiently co-immunoprecipitated with R4-SA, but showed weaker association with either wild-type RHBDL4 or the R4-SAUM construct (**Figs. 1D** and **EV1F-G**). ERGIC3, but not ERGIC2, additionally exhibited a ladder of higher molecular weight species characteristic of poly-ubiquitinated forms, which were diminished in R4-SAUM precipitates (**Fig. EV1G**). A similar pattern was observed for total immunoprecipitated ubiquitinated proteins (**Figs. 1D** and **EV1G**), consistent with other established RHBDL4 substrates (Fleig *et al*., 2012; Knopf *et al*., 2020). While the absence of detectable ubiquitinated ERGIC2 may reflect a distinct requirement for ubiquitination or technical limitations, these data suggest that RHBDL4 can engage trafficking regulators through distinct recognition modes, converging on proteolytic turnover as a common outcome. Taken together, these results support the identification of ERGIC2 and ERGIC3 as endogenous RHBDL4 substrates and indicate that their abundance is subject to RHBDL4-mediated quantity control.

### Molecular determinants of ERGIC recognition by RHBDL4

Having established the ERGIC family as a class of RHBDL4 substrates, we next sought to define the molecular determinants underlying their recognition and cleavage. Although RHBDL4 is classified as an intramembrane rhomboid protease, we previously demonstrated that it primarily cleaves ERAD substrates within their luminal domains (Fleig *et al*., 2012; Knopf *et al*., 2020). Consistent with this, cleavage site mapping using truncated constructs revealed luminal domain cleavage in all three ERGIC proteins (**Figs. 2A** and **EV2A-B**). AlphaFold-based structure predictions placed the cleavage sites in diverse secondary structures: β-sheet in ERGIC1, α-helices and loops/sheets in ERGIC2, and loops in ERGIC3 (**EV2C-E**), indicating that RHBDL4 can accommodate a wide range of structural contexts. Interestingly, the more efficiently cleaved sites align with a motif previously described for bacterial rhomboids and the plasma membrane protease RHBDL2 (Strisovsky *et al*, 2009). However, mutagenesis of this motif, as well as residues surrounding ERGIC2 residue 132, failed to abolish cleavage but rather enhanced processing (**Fig. EV2F-G**). Together with our previous observations for the RHBDL4 substrate MHC202 (Bock *et al*., 2022), these findings suggest that substrate recognition is not dictated by a strict consensus sequence but instead depends on higher-order structural features and substrate conformation.

**Figure 2.**
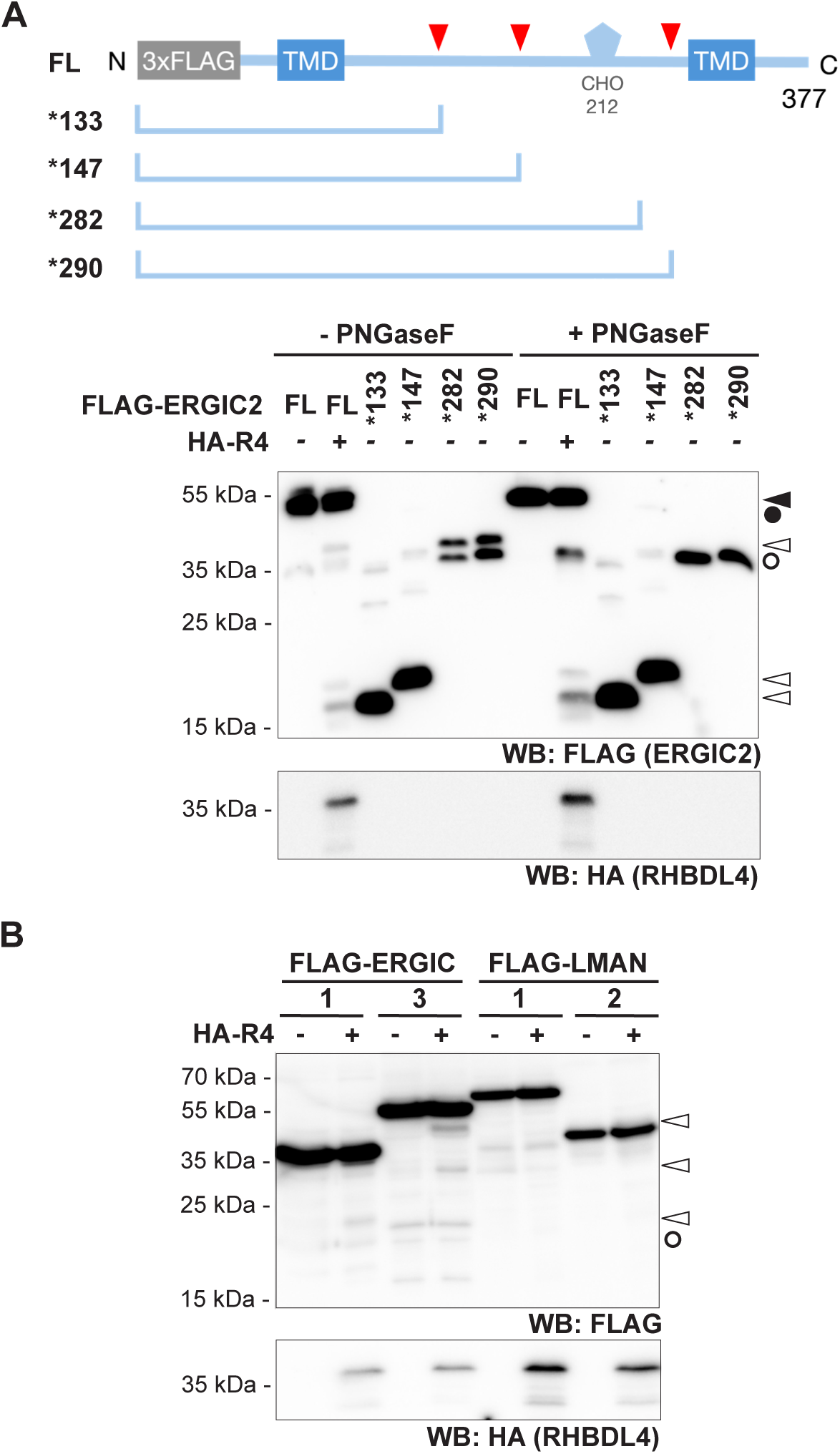
RHBDL4 cleaves ERGIC2 within its luminal domain. **(A)** Western blot (WB) analysis of HEK293T cells co-expressing full-length (FL) FLAG-ERGIC2 (closed arrow and circle) with either empty vector (-) or wild-type (WT) HA-RHBDL4 (HA-R4). FLAG-ERGIC2 truncations terminating after residues 133 (*133), 147 (*147), 282 (*282), and 290 (*290) were expressed and compared with the RHBDL4-generated cleavage fragments (open arrow and circle). Upper panel, schematic of the FLAG-ERGIC2 truncation mutants used as reference peptides. Comparison with the reference peptides identifies cleavage regions surrounding residues 132, 146, and 289, with the regions around residues 132 and 146 coinciding with predicted rhomboid cleavage motifs (see Fig. EV2C). Samples were left untreated or digested with PNGase F prior to analysis. Cells were treated with MG132 (2 µM) to stabilize cleavage fragments and reference peptides. **(B)** WB analysis comparing RHBDL4-dependent cleavage of FLAG-ERGIC1, FLAG-ERGIC3, FLAG-LMAN1, and FLAG-LMAN2 in HEK293T cells. Whereas ERGIC1 and ERGIC3 are efficiently cleaved by RHBDL4, LMAN1 and LMAN2 remain resistant to RHBDL4 processing. Cargo receptors were co-expressed either with an empty vector or WT HA-R4 and treated with MG132 (2 µM).

To further investigate RHBDL4 substrate specificity, we examined two members of the unrelated LMAN cargo receptor family, LMAN1/ERGIC-53 and LMAN2 (Satoh *et al*., 2014). In contrast to ERGIC proteins, neither receptor yielded prominent cleavage fragments, even when compared to the least efficiently cleaved ERGIC substrate (**Fig. 2C** and **EV3A-B**), indicating that cargo receptor identity alone is insufficient for RHBDL4 recognition. Previous work identified positively charged residues within TM domains as membrane-integral degrons that promote RHBDL4-dependent turnover (Fleig *et al*., 2012). To test whether such degrons contribute to cargo receptor recognition, we introduced a positively charged residue into the TM domain of LMAN2. Strikingly, the resulting I334K mutant became efficiently cleaved by RHBDL4 (**Fig. EV3C-D**), demonstrating that LMAN2 is intrinsically accessible to RHBDL4 once an appropriate degron is present. Conversely, mutation of a charge degron in the second TM domain of ERGIC2 significantly reduced cleavage efficiency (**Fig. EV3C** and **E**). Collectively, these findings indicate that RHBDL4 substrate recognition is governed by a combination of luminal substrate features and membrane-integral degrons. While TM degrons can strongly influence cleavage efficiency, they are not sufficient to explain substrate specificity, as ERGIC1 and ERGIC3 lack comparable motifs yet remain RHBDL4 substrates. These results support a model in which RHBDL4 selectively recognizes a subset of cargo receptors through multiple complementary determinants.

### RHBDL4 selectively tunes ERGIC-dependent trafficking pathways

ERGIC2 has been implicated in retrograde trafficking of the Wnt cargo receptor Evi/WLS (Yu *et al*., 2014). To test whether RHBDL4 modulates this pathway, we first examined its impact on the interaction between ERGIC2 and Evi. As HEK293T cells exhibit minimal endogenous Wnt activity, WNT3 was transiently expressed. Notably, shRNA-mediated knockdown of RHBDL4 increased the interaction between HA-tagged Evi and FLAG-tagged ERGIC2 (**Fig. 3A-B**), consistent with enhanced ERGIC2 availability upon loss of RHBDL4-mediated quantity control. HA-Evi levels were modestly elevated in input lysates from RHBDL4-depleted cells (**Fig. 3A**), prompting us to examine whether Evi itself is directly cleaved by RHBDL4. However, cleavage assays revealed negligible processing of Evi compared to FLAG-ERGIC2 (**Fig. EV4A-B**), suggesting Evi is not a primary RHBDL4 substrate, although indirect effects on Evi stability cannot be excluded. To determine whether this increased ERGIC2-Evi interaction has functional consequences, we stimulated Wnt signaling in HCT116 RHBDL4 knockout cells (**Fig. EV4C-D**) using the canonical Wnt ligand WNT3a. RHBDL4 knockout cells exhibited significantly increased expression of the Wnt target gene *AXIN2* (**Fig. 3C**), consistent with prior observations that ERGIC2 knockdown impairs Wnt signaling (Yu *et al*., 2014). Together, these findings are consistent with a role for RHBDL4 in limiting ERGIC2-dependent Wnt signaling.

**Figure 3.**
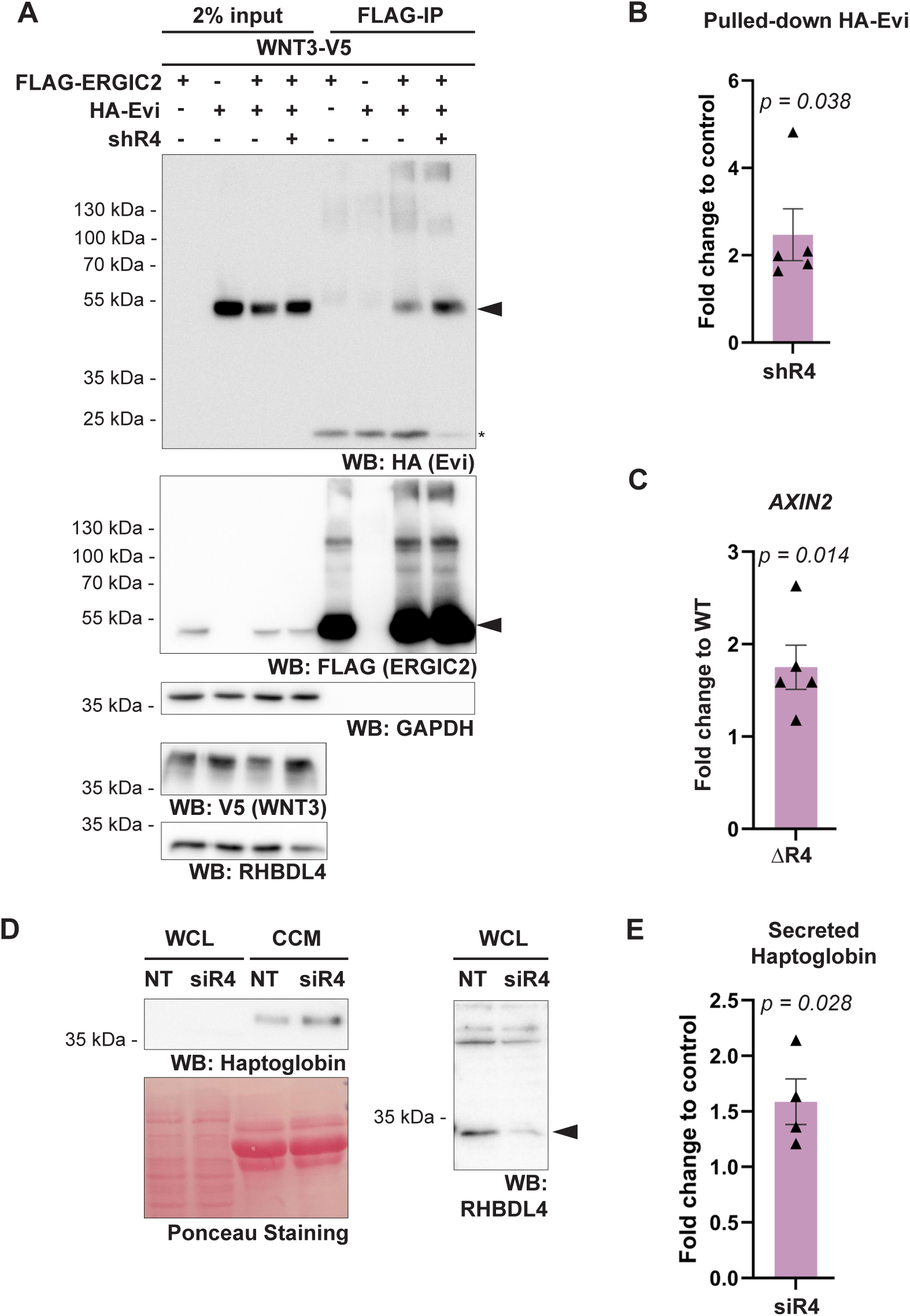
RHBDL4 regulates ERGIC-dependent trafficking pathways. **(A)** Western blot (WB) analysis of FLAG immunoprecipitations (IP) from HEK293T cells expressing either FLAG-ERGIC2, HA-Evi, or both together with WNT3-V5. RHBDL4 knockdown by shRNA (shR4) increases co-purification of HA-Evi with FLAG-ERGIC2 compared to control (NT). GAPDH serves as the loading control. **(B)** Quantification of the co-immunoprecipitation experiment shown in (A). Data represent means ± SEM (n = 5, *p < 0.05, unpaired two-sided Student’s t-test). **(C)** qRT-PCR analysis of *AXIN2* and *WNT3A* mRNA in HCT116 wild-type (WT) and RHBDL4 knockout (ΔR4) cells transiently expressing WNT3a. *AXIN2* expression is significantly increased in ΔR4 cells, indicating enhanced Wnt signaling. Data represents means ± SEM (n = 5, *p < 0.05; unpaired two-sided Student’s t-test). **(D)** WB analysis of secreted endogenous haptoglobin from HepG2 cells transfected with non-targeting (NT) or RHBDL4-targeting (siR4) siRNA. RHBDL4 knockdown increases haptoglobin secretion. Data represent means ± SEM, (n = 4, *p < 0.05; unpaired two-sided Student’s t-test). **(E)** Quantification of haptoglobin secretion shown in (D). Data represent means ± SEM (n = 4, *p < 0.05; unpaired two-sided Student’s t-test).

ERGIC3 has been identified as a cargo receptor for the secreted protein haptoglobin (Yoo *et al*., 2019), which is predominantly expressed in hepatocytes (Galicia & Ceuppens, 2011). To examine whether RHBDL4 also modulates ERGIC3-dependent trafficking, we assessed haptoglobin secretion in the hepatocyte-derived HepG2 cell line. siRNA-mediated knockdown of RHBDL4 significantly increased the secretion of endogenous haptoglobin (**Fig. 3D-E**) without elevating haptoglobin mRNA levels (**Fig. EV4E**). These findings indicate that RHBDL4 limits ERGIC3-dependent haptoglobin secretion at a post-transcriptional level, consistent with proteolytic quantity control of the cargo receptor.

Together, these data indicate that, in cultured cells, RHBDL4 influences at least two distinct ERGIC-dependent transport pathways: retrograde Evi trafficking and haptoglobin secretion. These findings support a broader role of RHBDL4 in selectively tuning secretory pathway dynamics through quantity control of ERGIC cargo receptors. Whether the observed phenotypes are mediated exclusively through turnover of ERGIC2 and ERGIC3 remains to be determined.

### RHBDL4 restrains secretion of lysosomal precursor proteins

Because RHBDL4 influenced multiple ERGIC-dependent trafficking pathways, we next asked whether its regulatory role extends more broadly to secretory pathway output. To address this, we performed an unbiased secretome analysis from wild-type and RHBDL4 knockout cells. Newly synthesized proteins were metabolically labeled with the methionine analog azidohomoalanine (AHA) for 24 hours, and secreted AHA-labeled proteins were enriched from conditioned medium via click chemistry, allowing stringent removal of serum-derived contaminants (**Fig. 4A**). Applying a cutoff of ±1.3-fold change and a p-value < 0.01, 69% of quantified secreted proteins remained unchanged, with a median fold change of 0.998, indicating preserved overall secretory capacity. Despite the absence of a global shift in secretion (**Fig. 4B**), several lysosomal proteins were significantly enriched in the RHBDL4 knockout secretome (**Table EV2**). Among these, cathepsin D (CatD), cathepsin L (CatL), PSAP, and progranulin (PGRN) showed increased secretion in the RHBDL4 knockout line used for the proteomic analysis. To validate these findings, we generated two additional independent RHBDL4 knockout clones (**Fig. EV5A**). Increased secretion of the pro-forms of CatD (pro-CatD), CatL (pro-CatL), and PSAP was reproducibly observed in all three RHBDL4 knockout clones (**Fig. 4C** and **EV5B-C**). PGRN secretion was likewise elevated in two of three knockout clones (**Fig. EV5D**), supporting the proteomic findings. Importantly, mRNA levels of CatD, CatL, and PSAP were largely unchanged in RHBDL4 knockout cells (**Fig. EV5E-G**), arguing that their enhanced secretion primarily reflects post-transcriptional regulation. However, PGRN mRNA levels were increased in RHBDL4-deficient cells (**Fig. EV5H**), indicating that transcriptional regulation may contribute to this phenotype. To validate our findings in an additional cell line, we extended our analysis to the lung fibroblast cell line WI-26-SV40 that has been extensively used to study subcellular trafficking (Nuchel *et al*, 2018; Nuchel *et al*, 2026; Nuchel *et al*, 2021) (**Fig. EV5I**). RHBDL4 depletion in WI-26 cells likewise resulted in increased secretion of pro-CatD and PSAP (**Fig. 4D**), while their mRNA levels remained unchanged (**Fig. EVJ**), supporting a post-transcriptional effect that is not restricted to HEK293T cells.

**Figure 4.**
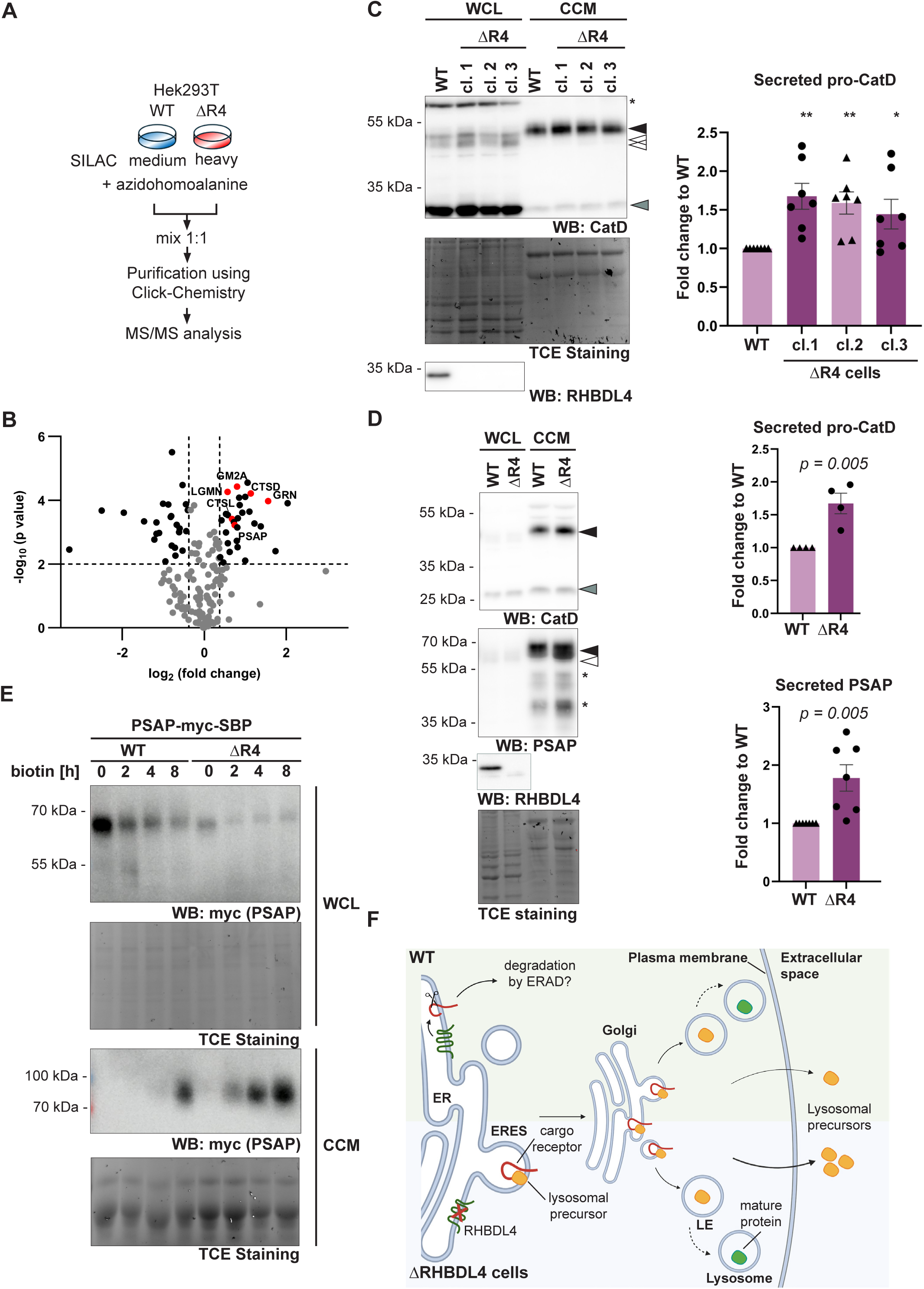
RHBDL4 restrains secretion of lysosomal precursor proteins. **(A)** Experimental outline of the mass-spectrometry-based secretome analysis in HEK293T wild-type (WT) and RHBDL4 knockout (ΔR4) cells. **(B)** Volcano plot of fold change and p-values for secreted proteins identified by quantitative secretome analysis (n = 4). Lysosomal proteins are highlighted in red. **(C)** Western blot (WB) analysis of endogenous CatD secretion in HEK293T WT and ΔR4 cells. Secretion of the CatD pro-form (closed arrow) is significantly increased in all three RHBDL4 knockout clones. The intracellular pro-form migrates at a lower apparent molecular weight (open arrow) whereas the mature heavy chain is detected mainly in the cell lysates (grey arrow). Background bands are marked with an asterisk. TCE fluorescence serves as the loading control (means ± SEM, n = 4, *p < 0.05, **p < 0.01, unpaired two-sided Student’s t-test). **(D)** WB analysis of CatD and PSAP secretion in WI-26 cells (means ± SEM, n = 4 (CatD), n = 7 (PSAP), **p < 0.01, unpaired two-sided Student’s t-test). **(E)** WB analysis of PSAP-myc-SBP trafficking using the RUSH assay in WT and ΔR4 WI-26 cells. Following biotin addition, intracellular PSAP-myc-SBP is depleted more rapidly and secreted more efficiently in ΔR4 cells (n = 3). **(F)** Working model of RHBDL4-dependent regulation of lysosomal precursor secretion. RHBDL4 limits precursor secretion by regulating trafficking decisions upstream of lysosomal delivery without affecting canonical M6P-dependent lysosomal targeting or lysosomal maturation.

Lysosomal proteins are typically synthesized as inactive precursor forms that undergo proteolytic maturation upon reaching the acidic environment of the lysosome (Mach, 2002). Under certain conditions, however, precursor forms can also be secreted, although the physiological relevance of this process remains incompletely understood (Benes *et al*, 2008; Carvelli *et al*, 2015; Ishidoh & Kominami, 1998). Notably, secreted pro-CatD migrated slightly slower than its intracellular precursor on SDS-PAGE, possibly reflecting differences in the post-translational modification. The mature CatD heavy chain (30 kDa), which results from an autocatalytic cleavage producing both heavy and light chains (Zaidi *et al*, 2008), were not decreased intracellularly, indicating that lysosomal delivery was not impaired (**Fig. EV6A**). Similar observations were made for the single chain of mature Cathepsin L (**Fig. EV6B**). In addition, the number of lysosomes per cell, assessed by LysoTracker staining, was unchanged in RHBDL4 knockout cells (**Fig. EV6C**), further indicating that lysosomal function remained intact. Consistent with the increased secretion of PSAP in HEK293T (**Fig. EV5B**) and WI-26 ΔR4 cells (**Fig. 4D**), analysis of PSAP trafficking using the RUSH system, which enables synchronized release of cargo from the ER (Boncompain *et al*, 2012), further supported enhanced PSAP secretion in RHBDL4-deficient cells (**Fig. 4E**). Together, these data indicate that RHBDL4 loss selectively enhances secretion of lysosomal precursor proteins by increasing their trafficking through the secretory pathway, without impairing intracellular lysosomal maturation.

### Increased lysosomal precursor secretion is not driven by direct substrate cleavage or impaired M6P trafficking

To explore the mechanism underlying increased secretion of lysosomal precursor proteins in RHBDL4 knockout cells, we first examined whether these proteins are direct RHBDL4 substrates. Co-expression cleavage assays showed minimal processing of pro-CatD and pro-CatL by RHBDL4 (**Fig. EV6D-E**), arguing against direct cleavage as a regulatory mechanism.

In contrast, a prominent RHBDL4-dependent cleavage fragment was observed for PSAP (**Fig. EV6F**), consistent with its prior identification in an RHBDL4 substrate screen (Tang *et al*, 2022). However, this fragment was not stabilized by proteasome inhibition (**Fig. EV6F-G**), indicating that RHBDL4-mediated PSAP cleavage is functionally distinct from ERAD.

These findings suggest that the increased secretion phenotype in RHBDL4-deficient cells does not arise from direct precursor cleavage or defective lysosomal maturation but rather reflects altered trafficking fates within the secretory pathway. We therefore next examined whether M6P-dependent lysosomal targeting was affected. However, cellular M6P levels were unchanged in RHBDL4 knockout cells (**Fig. EV6H**), and together with the finding that mature lysosomal forms of CatD and CatL were largely unaltered (**Fig. EV6A-B**), indicating that the canonical M6P pathway remains intact. Instead, these data suggest that in the absence of RHBDL4, a larger fraction of lysosomal precursors exit the ER and is diverted into the secretory pathway. Together, these findings argue that RHBDL4 regulates lysosomal precursor secretion indirectly through one or more trafficking regulators rather than through direct substrate cleavage or disruption of canonical lysosomal targeting pathways (**Fig. 4F**).

### Sortilin is an RHBDL4 substrate that promotes non-canonical lysosomal precursor secretion

Based on these observations, we hypothesized that RHBDL4 limits non-canonical lysosomal precursor secretion by controlling the abundance of such an alternative trafficking regulator. One such regulator is the lysosomal sorting receptor sortilin, which emerged from a previous proteomic substrate screen for RHBDL4 (Knopf *et al*., 2020). Sortilin functions as a lysosomal cargo receptor that acts independently of the M6P pathway and has been implicated in the trafficking of both PSAP (Hassan *et al*, 2004; Lefrancois *et al*., 2003) and CatD (Canuel *et al*., 2008).

To test whether sortilin is a direct RHBDL4 substrate, we performed a rhomboid cleavage assay. Co-expression of sortilin with wild-type RHBDL4 yielded a ∼20 kDa cleavage fragment, which was stabilized upon proteasome inhibition with MG132, consistent with RHBDL4-dependent ERAD (**Fig. 5A**). In addition, co-expression with catalytically inactive RHBDL4 led to accumulation of deglycosylated full-length sortilin (**Fig. 5A**), a hallmark of retrotranslocation intermediates previously observed for other RHBDL4 substrates (**Fig. EV1A**, Bock *et al*., 2022). Based on its size, the RHBDL4-generated fragment likely separates the luminal cargo-binding domain of sortilin from its cytosolic tail. Such cleavage would be predicted to uncouple cargo recognition from clathrin adaptor recruitment function of sortilin, thereby rendering the resulting fragments functionally inactive (**Fig. 5A**). In line with our findings, ubiquitin-dependent cleavage of the sortilin homolog Vps10 by the related rhomboid protease Rbd2 has been reported in *Saccharomyces cerevisiae* (Minard *et al*, 2026).

**Figure 5.**
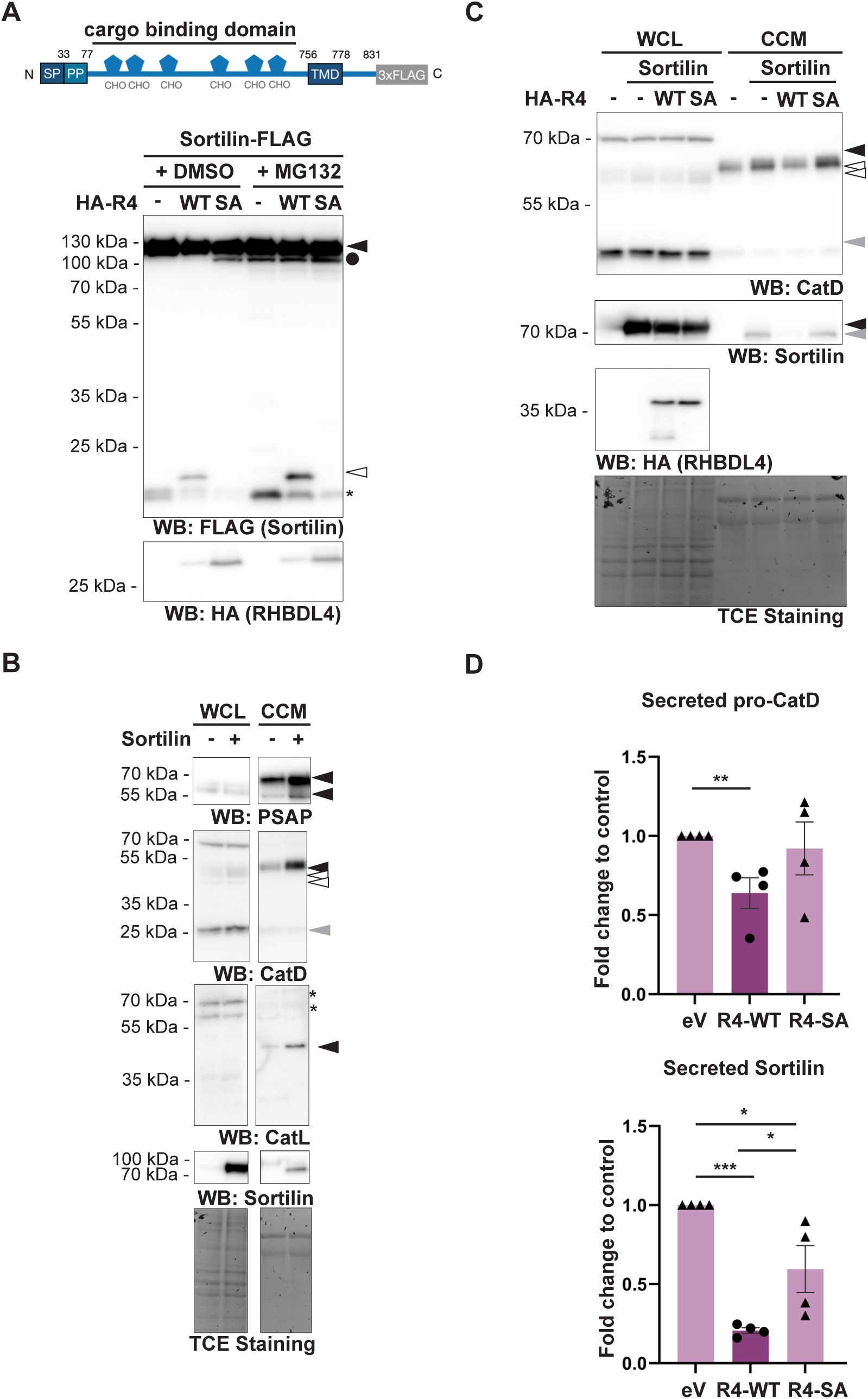
RHBDL4 antagonizes sortilin-dependent lysosomal precursor protein secretion. **(A)** Western blot (WB) analysis of sortilin-FLAG (closed arrow) ectopically expressed in HEK293T cells together with empty vector (-), wild-type (WT) HA-RHBDL4 (HA-R4), or the catalytic SA mutant. WT HA-R4 generates an N-terminal FLAG-sortilin cleavage fragment (open arrow). Cells were treated with either the DMSO or MG132 (2 µM). TCE fluorescence serves as the loading control. Asterisk indicates nonspecific bands. Upper panel, schematic of the sortilin-FLAG construct; SP, signal peptide; PP, pro-peptide; CHO, N-linked glycosylation site. **(B)** WB analysis of PSAP, CatD, and CatL secretion in HEK293T cells expressing ectopic sortilin. sortilin expression increases secretion of all three lysosomal precursor proteins. TCE fluorescence serves as the loading control. **(C)** WB analysis of CatD secretion in HEK293T cells co-expressing sortilin with empty vector, WT HA-R4, or the catalytic SA mutant. WT HA-R4 suppresses sortilin-dependent CatD secretion (closed arrow) and reduces secretion of sortilin itself (grey arrow). TCE, loading control. **(D)** Quantification of secretion assay shown in (C). Data represents means ± SEM (n = 4, *p < 0.05, **p < 0.01, unpaired two-sided Student’s t-test).

To determine whether elevated sortilin levels are sufficient to drive increased lysosomal precursor secretion, we overexpressed sortilin in HEK293T cells. This resulted in increased secretion of pro-CatD, pro-CatL, and PSAP (**Fig. 5B**), closely mimicking the RHBDL4 ablation phenotype. Importantly, levels of mature intracellular CatD and CatL remained largely unchanged (**Fig. 5B**), indicating that under these conditions sortilin preferentially promotes secretion of unprocessed precursor forms rather than disrupting lysosomal delivery. Consistent with previous reports, sortilin itself was detected in the culture medium as a faster migrating species relative to its intracellular form, indicative of ectodomain shedding (Hermey *et al*, 2006; Nyborg *et al*, 2006). Consistently, inhibition of ADAM metalloproteases with BB94 caused a shift to a higher molecular weight form, further supporting ectodomain shedding as the source of secreted sortilin (**Fig. EV7**). To test whether RHBDL4 can modulate sortilin-induced precursor secretion, we co-expressed sortilin with either wild-type or catalytically inactive RHBDL4. Only wild-type RHBDL4 suppressed the sortilin-driven increase in secretion of pro-CatD **(Fig. 5C-D**), demonstrating that RHBDL4 antagonizes sortilin in a protease activity-dependent manner. Notably, RHBDL4 co-expression also reduced the amount of secreted sortilin (**Fig. 5C-D**), suggesting that RHBDL4 limits the pool of sortilin that reaches late secretory compartments where ectodomain shedding occurs (Lichtenthaler *et al*, 2018).

In summary, these results indicate that RHBDL4 counteracts sortilin-dependent secretion of lysosomal precursors by limiting sortilin functional abundance. While this mechanistic model is currently based largely on overexpression experiments, it provides initial evidence that RHBDL4 regulates non-canonical lysosomal cargo trafficking through quantity control of a key trafficking receptor.

## Discussion

Our study identifies cargo receptors as a major functional class of RHBDL4 substrates and establishes RHBDL4 as a regulator of secretory pathway organization. Rather than directly controlling individual cargo proteins, RHBDL4 remodels trafficking flux by proteolytically tuning cargo receptor abundance, thereby regulating distinct transport pathways, including ERGIC-dependent trafficking and sortilin-mediated lysosomal precursor secretion.

### RHBDL4 selectively targets cargo receptors to regulate trafficking flux

Selective regulation of protein trafficking requires RHBDL4 to recognize cargo receptors with defined substrate specificity. Consistent with previous work, RHBDL4 substrate recognition depends on a combination of TM domain features, ubiquitination, and local structural permissiveness at the cleavage site rather than a strict consensus sequence (Bock *et al*., 2022; Fleig *et al*., 2012; Knopf *et al*., 2020). Accordingly, ERGIC cargo receptors are efficiently cleaved despite substantial sequence divergence at their cleavage sites, whereas related LMAN proteins containing apparent consensus motifs are not processed.

Our data further indicate that RHBDL4 employs distinct substrate recognition modes. Whereas ERGIC3 interacts with RHBDL4 in a ubiquitin-dependent manner, similar to TMED7 (Knopf *et al*., 2024), ERGIC2 appears to be recognized independently of detectable ubiquitination, likely through a charged TM degron. Together with the observation that ERGIC3 is regulated by the ubiquitin ligase MARCH2 (Yoo *et al*., 2019), these findings support a model in which RHBDL4 cooperates with ubiquitin-dependent quality control pathways to regulate the abundance of selected trafficking regulators.

Collectively, our data suggest that cargo receptors constitute a selective class of RHBDL4 substrates. While TMED7, ERGIC2, ERGIC3, and sortilin are efficiently cleaved, related cargo receptors such as LMAN1, LMAN2, and several other TMED family members remain refractory to RHBDL4 processing (Knopf *et al*., 2024). This selectivity indicates that RHBDL4 preferentially targets specific key trafficking determinants rather than broadly controlling cargo receptor turnover. By acting one hierarchical level upstream of individual cargos, RHBDL4 remodels secretory pathway flux through selective control of cargo receptor abundance.

### RHBDL4 remodels ERGIC-dependent transport pathways

Consistent with this model, RHBDL4 negatively regulates canonical Wnt signaling in HCT116 cells, most likely by limiting ERGIC2-dependent trafficking of the Wnt cargo receptor Evi. Although the physiological relevance remains to be established, these findings are consistent with previous reports linking RHBDL4 to Wnt signaling (Penalva *et al*, 2025). Given that RHBDL4 is upregulated in colorectal cancer, where Wnt signaling is constitutively activated (Zhang *et al*, 2018), RHBDL4 induction may represent a negative feedback mechanism that restrains excessive pathway activation, analogous to its previously described role in limiting TLR4 signaling through TMED7 cleavage (Knopf *et al*., 2024).

We further identify ERGIC3 as an endogenous RHBDL4 substrate and show that RHBDL4 modulates ERGIC3-dependent secretion of haptoglobin in HepG2 cells (Yoo *et al*., 2019).

Although also here the physiological consequences remain to be established, this finding further illustrates how proteolytic regulation of cargo receptor abundance can selectively reshape secretory output without globally affecting protein secretion.

### RHBDL4 restricts sortilin-dependent secretion of lysosomal precursor proteins

Our unbiased secretome analysis identified pro-CatD, pro-CatL, and PSAP as secretory cargos negatively regulated by RHBDL4. Mechanistically, our data identify the lysosomal cargo receptor sortilin as a central mediator of this pathway. Although sortilin is well established as an alternative lysosomal sorting receptor for cargos such as PSAP and CatD (Canuel *et al*., 2008; Hassan *et al*., 2004; Lefrancois *et al*., 2003; Zeng *et al*, 2009), its contribution to protein secretion has remained less clear.

Our findings demonstrate that increased sortilin abundance is sufficient to enhance secretion of lysosomal precursor proteins, identifying a previously unappreciated secretory function of sortilin. RHBDL4 attenuates this pathway by proteolytically limiting a trafficking-competent pool of sortilin, thereby reducing secretion of pro-CatD while leaving canonical M6P-dependent lysosomal targeting largely unaffected. The absence of detectable changes in total cellular sortilin levels suggests that RHBDL4 selectively regulates a functionally distinct receptor pool rather than the entire cellular population. This interpretation is consistent with previous observations that oxidative stress redirects PSAP secretion through disruption of sortilin-mediated trafficking (Toyofuku *et al*, 2012), supporting the idea that sortilin acts as a context-dependent determinant of trafficking fate by balancing lysosomal delivery and secretion.

The physiological significance of regulated secretion of lysosomal precursor proteins remains incompletely understood. However, several of the proteins identified here possess well-established extracellular functions, including a neuroprotective role of PSAP, tumor-promoting activities of secreted pro-CatD and pro-CatL, and immunomodulation by progranulin (Baker *et al*, 2006; Barbarin & Frade, 2011; Benes *et al*., 2008; He *et al*, 2003; Lan *et al*, 2021; O’Brien *et al*, 1994; Zhu *et al*, 2023). These observations suggest that RHBDL4-dependent regulation of lysosomal precursor secretion may influence physiological processes extending well beyond intracellular lysosome biology.

Our findings therefore establish RHBDL4 as a proteolytic rheostat that regulates secretory pathway architecture through selective turnover of cargo receptors. Rather than functioning only as a quality-control protease that eliminates aberrant proteins, RHBDL4 dynamically adjusts the abundance of trafficking determinants to remodel transport capacity while preserving overall secretory competence. Such hierarchical regulation provides an efficient mechanism for coordinating diverse secretory programs and may become particularly important during cellular stress or immune activation, conditions in which RHBDL4 expression is induced (Knopf *et al*., 2024). This model also provides a conceptual framework for understanding why RHBDL4 deficiency causes relatively modest phenotypes under basal conditions while permitting context-dependent remodeling of secretory pathway function (Lastun *et al*, 2022).

### Limitations of the study

While our findings identify cargo receptor quantity control as a central function of RHBDL4, several important questions remain. In particular, the physiological signals that dynamically regulate RHBDL4-dependent cargo receptor turnover and the extent to which this mechanism is engaged across different tissues and stress conditions remain to be determined. In addition, although our data establish direct RHBDL4-dependent cleavage of several cargo receptors, the broader spectrum of regulated trafficking pathways and their physiological relevance will require further investigation.

## Methods

### Plasmids

All constructs were cloned into pcDNA3.1(+) (Invitrogen). Constructs encoding human RHBDL4 and its catalytic S144A mutant, carrying either an N-terminal triple HA-tag or a C-terminal GFP-tag, have been described previously (Fleig *et al*., 2012). Human ERGIC1 (Gene ID 57222) was generated by introducing the V146I and T149V substitutions into the murine sequence (Gene ID 67458, full-ORF Gateway cDNA clone, GenBank accession CT010230) and subcloned into a pcDNA3.1-based expression vector containing an N-terminal triple FLAG-tag. Open reading frames encoding ERGIC2 (Gene ID 51290) and ERGIC2-Ala7 were synthesized as custom gene fragments (Integrated DNA Technologies) and subcloned into the same vector. ERGIC3 (Gene ID 51614, full-ORF Gateway cDNA clone, GenBank accession HQ448277) and sortilin (Gene ID 6272, full-ORF Gateway cDNA clone, GenBank accession DQ896793) were amplified by PCR and subcloned into pcDNA3.1-based expression vectors containing either an N-terminal triple FLAG-tag or, in the case of sortilin, a C-terminal triple FLAG-tag. Open reading frames encoding human Evi/WLS (Gene ID 79971), CatL (Gene ID 1514), CatD (Gene ID 1509), PSAP (Gene ID 5660), and WNT3a (Gene ID 89780) were amplified from HEK293T cDNA by PCR. Evi was subcloned into pcDNA3.1-based expression vectors containing either an N-terminal triple FLAG-tag or an N-terminal triple HA-tag. PSAP, CatD, and CatL were cloned with a C-terminal triple FLAG-tag, whereas WNT3a was cloned without an epitope-tag. Human LMAN1 (Gene ID 3998) was amplified from HEK293T cDNA without its signal peptide, whereas human LMAN2 (Gene ID 10960) was amplified from WI-26 cDNA. Both coding sequences were subcloned into a pcDNA3.1-based expression vector containing a signal peptide upstream of an N-terminal triple FLAG-tag. Cleavage-site point mutants, TM domain charge mutants, reference peptides, and an untagged sortilin construct were generated by site-directed mutagenesis. For the RUSH assay, the previously published pIRES-Str-KDEL-hPSAP-SBP construct (Nuchel *et al*., 2026) was modified by in-frame insertion of a Myc-tag sequence between the PSAP cDNA and the SBP-tag (PSAP-RUSH).

### Culture and generation of cell lines

HEK293T wild-type cells (CRL-3216, ATCC), HEK293T RHBDL4 knockout cells, Hek293T GNPTAB knockout cells (Fernandes *et al*, 2024) and HEK293T ERGIC2-3xFLAG cells were cultured in DMEM (Gibco) supplemented with 10% fetal bovine serum (FBS) at 37°C in a humidified atmosphere containing 5% CO₂. HCT116 cells (gift from Elmar Schiebel, ZMBH, Heidelberg, Germany) were cultured in McCoy’s 5A medium (Gibco) supplemented with 10% FBS. HepG2 cells (gift from Bernhard Dobberstein, ZMBH, Heidelberg, Germany) were cultured in MEM (Gibco) supplemented with 10% FBS. WI-26 SV40 fibroblasts (WI-26; CCL-95.1, ATCC) were cultured in DMEM/F12 GlutaMAX medium (Gibco) supplemented with 10% FBS. Unless otherwise indicated, all cells were maintained at 37°C in a humidified atmosphere containing 5% CO₂. RHBDL4 knockout clone 1 (#81), generated using TALEN expression vectors, has been described previously (Knopf *et al*., 2020). RHBDL4 knockout clone 2 (2B6) and clone 3 (2G9) were generated by transfecting HEK293T cells with a plasmid encoding Cas9 and a single guide RNA targeting RHBDL4 (forward: 5′-caccgGTTGAGGGCCAAAGTTGCTA-3′; reverse: 5′-aaacTAGCAACTTTGGCCCTCAACc-3′). Transfected cells were selected with puromycin (1 μg/ml) for 7 days. Single-cell clones were isolated by limiting dilution and validated by western blotting and genomic DNA sequencing. Specifically, the RHBDL4 locus was amplified from genomic DNA using the following primers: forward, 5′-AGCATTTGCCTCATGGGAGT-3′; reverse, 5′-CAAGACGCAAAGCCTCAGC-3′. Editing outcomes were analyzed using TIDER (Brinkman *et al*, 2018). HCT116 RHBDL4 knockout cells were generated using the same strategy. Endogenously tagged ERGIC2-3xFLAG HEK293T cells were generated by CRISPR/Cas12-mediated genome editing (Fueller *et al*, 2020). The tagging cassette was amplified by PCR using primers containing homology arms targeting the ERGIC2 locus (M1: 5′-TGGTCTTTTTTTCCCTCCTTTTCTCCTTAGGTTCCTTTTGAGGATGGCCACACAGACAACCACTTACCTCTTTTAGAAAATAATACACATTCAGGTGGAGGAGGTAGTG-3′; M2: 5′-TAAAAAAAGGTTTTATGTCTCAGGCAAAAAGTTTTTCTCCTTCAATCGGGAGGTGAAAAAAACACATTAACACCTCCCGATATCTACACTTAGTAGAAATTAGCTAGCTGCATCGGTAC-3′). Cells were co-transfected with the tagging cassette and a Cas12a expression plasmid. Positive cells were selected with zeocin (500 ng/ml) for 5 days. Single-cell clones were isolated by limiting dilution and validated by western blotting and genomic DNA sequencing. WI-26 RHBDL4 knockout cells were generated using CRISPR/Cas9 (Ran *et al*, 2013) with double-stranded DNA oligonucleotides targeting RHBDL4 (5′-caccgCAAGCAAATCTGTTCGGTAC-3′ and 5′-aaacGTACCGAACAGATTTGCTTGC-3′) as described previously (Nuchel *et al*., 2021). Cells were transfected with the Cas9 expression plasmid and guide RNA construct, selected with zeocin (2 μg/ml) for 5 days, and single-cell clones were isolated by limiting dilution and validated by western blotting and genomic DNA sequencing.

### Transfection

For ectopic protein expression, cells were transfected with plasmid DNA 24 h after seeding using 25 kDa linear polyethyleneimine (Polysciences) (Durocher *et al*, 2002). Unless otherwise indicated, cleavage assays were performed by co-transfecting 500 ng of substrate construct with 200 ng of RHBDL4 expression construct in 6-well plates. For substrate-trapping experiments, 1.8 μg of RHBDL4 expression construct was transfected into cells cultured in 10 cm dishes. FLAG immunoprecipitation experiments were performed by transfecting 3 μg of ERGIC2, Evi, or WNT3 expression constructs per 10 cm dish. shRNA-mediated knockdown of RHBDL4 was achieved by transfecting 4.8 μg of plasmid encoding either RHBDL4-targeting or control shRNA. Empty vector DNA was added where necessary to keep the total amount of transfected DNA constant across all conditions. For siRNA-mediated knockdown, cells were transfected in 12-well plates with 50 pmol of RHBDL4-targeting siRNA (ON-TARGETplus SMARTpool Human siRNA, Horizon Discovery, L-019378-00-0005; sequences: 5′-CGGCAAUACUACUUUAAUA-3′, 5′-CGAGGAAAUACCAGAAAUA-3′, 5′-GGGAUAAAUACUGGACUUA-3′, 5′-GACAGCGGCUUCACAGAUU-3′) or non-targeting siRNA (D-001810-10-20; sequences: 5′-UGGUUUACAUGUCGACUAA-3′, 5′-UGGUUUACAUGUUGUGUGA-3′, 5′-UGGUUUACAUGUUUUCUGA-3′, 5′-UGGUUUACAUGUUUUCCUA-3′) using Lipofectamine RNAiMAX (Thermo Fisher Scientific). Proteasomal degradation was inhibited by treatment with 2 μM MG132 (Calbiochem) for 16 h. p97/VCP was inhibited with CB-5083 (ApexBio) at 1 μM for 16 h in cleavage assays or at 2.5 μM for 5 h prior in immunoprecipitation experiments.

### Immunoprecipitation

Cells were lysed for 60 min on ice in solubilization buffer (50 mM HEPES-KOH, pH 7.4, 150 mM NaCl, 2 mM MgOAc₂, 10% glycerol, 1 mM EGTA, and 10 mM N-ethylmaleimide) supplemented with 1% Triton X-100 (Merck) and EDTA-free Complete protease inhibitor cocktail (Roche). Triton X 100-insoluble material was removed by centrifugation at 16,000 x g for 15 min at 4°C. The supernatants were diluted 1:1 with detergent-free lysis buffer and precleared with Protein G Sepharose beads (GE Healthcare) for 2 h. For ERGIC3 trapping immunoprecipitations, GFP-nanobody beads were added, whereas FLAG immunoprecipitations were performed using anti-FLAG M2 agarose (Sigma). For ERGIC2-FLAG trapping immunoprecipitations, the preclearing step was omitted and GFP antibody (1.6 μg; 11814460001, Roche) was added directly. Samples were incubated overnight at 4°C with gentle rotation. Protein G magnetic beads (Thermo Fisher Scientific) were subsequently added to the ERGIC2-FLAG trapping immunoprecipitations for 1 h. Beads were washed either four times with lysis buffer (ERGIC3 trapping and FLAG immunoprecipitations) or four times with wash buffer (TBS containing 0.5% Tween-20, 0.5 M NaCl, EDTA-free complete protease inhibitor) (ERGIC2-FLAG trapping immunoprecipitations), before bound proteins were eluted in SDS sample buffer (50 mM Tris, pH 6.8, 10 mM EDTA, pH 8.0, 3.75% glycerol, 2% SDS, 0.01% bromophenol blue, and 5% β-mercaptoethanol). Samples were analyzed by western blotting.

### Cellular fractionation and proteomic analysis

For the RHBDL4 substrate screen, two biological replicates were analyzed. HEK293T wild-type or RHBDL4 knockout cells (Knopf *et al*., 2020) were SILAC-labeled for approximately two weeks with either unlabeled (light) or heavy (146 μg/ml Lys-8, 84 μg/ml Arg-10) amino acids (Silantes). Twenty-four hours before harvesting, cells of each genotype were seeded into 10 cm dishes. Cells were washed, detached, combined, and pelleted by centrifugation at 500 x g for 5 min at 4°C. Pellets were resuspended on ice in hypotonic buffer (10 mM HEPES-KOH, pH 7.4, 1.5 mM MgCl_2_, 10 mM KCl) supplemented immediately before use with PMSF (10 μg/ml), 0.5 mM DTT, and EDTA-free Complete protease inhibitor cocktail (Roche). Following 10 min of swelling on ice, cells were lysed by passing the suspension five times through a 27-gauge needle. Cell debris and nuclei were removed by centrifugation at 1,000 x g for 10 min at 4°C. Membranes were collected from the resulting supernatant by centrifugation at 100,000 x g for 20 min and resuspended in rough microsome buffer (50 mM HEPES-KOH, pH 7.4, 250 mM sucrose, 50 mM KOAc, 2 mM Mg(OAc)_2_) supplemented with 1 mM DTT. KOAc and EDTA concentrations were then adjusted to 500 mM and 50 mM, respectively, and samples were incubated on ice for 10 min. Membranes were layered onto a high-salt sucrose cushion (50 mM HEPES-KOH, pH 7.4, 500 mM sucrose, 500 mM KOAc, 5 mM Mg(OAc)_2_) and centrifuged at 140,000 x g for 30 min. The resulting pellet was resuspended in 50 μl SDS sample buffer (50 mM Tris, pH 6.8, 10 mM EDTA, pH 8.0, 3.75% glycerol, 2% SDS, 0.01% bromophenol blue, 5% β-mercaptoethanol) and separated by SDS-PAGE on a 4-12% Novex NuPAGE Bis-Tris gel (Thermo Fisher Scientific) until proteins had migrated approximately 1 cm into the resolving gel. Gels were stained with Quick Coomassie Stain (Serva), cut into two pieces, and washed with 50% acetonitrile. Proteins were reduced with DTT, alkylated with iodoacetamide, and digested with trypsin in 0.01% trifluoroacetic acid. Peptides were extracted with 50% acetonitrile and 10% formic acid, acidified with 1% trifluoroacetic acid, and submitted to the Core Facility for Mass Spectrometry and Proteomics at the Centre for Molecular Biology Heidelberg for LC-MS/MS analysis as described previously (Knopf *et al*., 2024).

### Secretome analysis

For quantitative secretome analysis, four biological replicates of HEK293T RHBDL4 knockout cells were analyzed. Cells were SILAC-labeled for two weeks with either heavy (146 μg/ml Lys-8, 84 μg/ml Arg-10) or medium (146 μg/ml Lys-4, 84 μg/ml Arg-6) amino acids (Silantes). Twenty-four hours before harvesting, cells were incubated for 30 min in methionine-free SILAC medium (Enzo Life Sciences) supplemented with 10% dialyzed FBS (Silantes), followed by the addition of methionine-free SILAC medium containing 100 μM azidohomoalanine (AHA; 63669AS, Anaspec). Conditioned media from wild-type and RHBDL4 knockout cells were combined, cleared of dead cells by centrifugation, and concentrated using 3 kDa molecular weight cut-off centrifugal filters (Merck). AHA-labeled proteins were captured using the Click Chemistry Capture Kit (CLK-1065, Jena Biosciences) according to the manufacturer’s instructions by coupling to alkyne agarose (CLK-1032-2, Jena Biosciences) at room temperature overnight. Samples were reduced with 5 mM DTT and alkylated with 40 mM chloroacetamide. To remove nonspecifically bound proteins and serum components, the beads were transferred to columns (Jena Biosciences) and washed sequentially with 20 ml each of SDS wash buffer (Click Chemistry Capture Kit), 8 M urea in 100 mM Tris-HCl (pH 8.0), 20% isopropanol, and 20% acetonitrile. Proteins were digested on-bead with 1 μg trypsin and 0.5 μg Lys-C (37286, Serva) at 37°C and 900 rpm in a shaker overnight. Samples were acidified with 10% formic acid and purified using SDB-RP StageTips prior to mass spectrometric analysis. Samples were analyzed at the CECAD Proteomics Facility using an Orbitrap Exploris 480 mass spectrometer equipped with a FAIMS Pro interface and coupled to an UltiMate 3000 nanoLC system (Thermo Fisher Scientific). Peptides were loaded onto a PepMap trap cartridge (#160434, Thermo Fisher Scientific) using solvent A (0.1% formic acid in water) and reverse-flushed onto an in-house packed analytical column (50 cm x 75 μm inner diameter, packed with 2.7 μm Poroshell EC120 C18 resin; Agilent). Peptides were separated at a constant flow rate of 300 nl/min using the following gradient: 3% solvent B (0.1% formic acid in 80% acetonitrile) to 5% within 1 min, 30% within 107 min, and 50% over the subsequent 20 min, followed by washing at 95% solvent B and column equilibration. The mass spectrometer was operated in data-dependent acquisition mode using three rotating FAIMS compensation voltages (-45, -60, and -75 V). For each compensation voltage, MS1 survey scans were acquired over an m/z range of 350-1,400 at a resolution of 60,000. The 12 most abundant precursor ions were isolated using a 1.4 Th isolation window and fragmented by higher-energy collisional dissociation (HCD) at a normalized collision energy of 30%. The AGC target was set to 100% with a maximum injection time of 60 ms. Fragment ions were detected in the Orbitrap at a resolution of 30,000. Dynamic exclusion was set to 25 s. Raw data were first separated into individual FAIMS compensation voltage files using FreeStyle (Thermo Fisher Scientific) before processing with MaxQuant (version 2.2) [10.1038/nprot.2016.136]. Database searches were performed against the UniProt canonical human proteome (UP5640; downloaded 20 January 2023) using default parameters with the match between runs option enabled across biological replicates. Raw files corresponding to the three compensation voltages of each sample were assigned identical sample names to allow their reintegration into a single output. Downstream statistical analysis was performed in Perseus (version 1.6.15) [10.1038/nmeth.3901]. Reverse database hits, contaminants, and proteins identified only by modified peptides were removed. Data were filtered for completeness across replicate groups, missing LFQ values were imputed using the default sigma-downshift approach, and differential abundance was assessed using FDR-controlled t-tests with s₀ = 0.2.

### Antibodies

The following antibodies were used for western blotting with indicated dilutions: 1:1000 rabbit polyclonal anti-calnexin (ab22595, Abcam), 1:1000 mouse monoclonal anti-FLAG M2 (F1804, Sigma Aldrich), 1:1000 mouse monoclonal anti-FLAG-HRP (clone M2, A8592, Sigma Aldrich), 1:1000 rabbit polyclonal anti-RHBDL4 (HPA013448, Sigma), 1:1000 rat monoclonal anti-HA (clone 3F10, 11867431001, Roche), 1:3000 rabbit monoclonal anti-ERGIC3 (clone EPR8141, ab129179, Abcam), 1:1000 rabbit polyclonal anti-haptoglobin (16665-1-AP, Proteintech), 1:1000 mouse monoclonal anti-myc (clone 9B11, #2276, Cell Signaling Technologies) 1:2000 rabbit polyclonal anti-PSAP (10801-1-AP, Proteintech), 1:2000 rabbit polyclonal anti-CatL (27952-1-AP, Proteintech), 1:2000 rabbit polyclonal anti-CatD (21327-1-AP, Proteintech), 1:1000 rabbit polyclonal anti-sortilin (12369-1-AP, Proteintech), 1:1000 mouse monoclonal anti-Ubiquitin (clone P4D1, sc-8017, Santa Cruz Biotechnology), 1:1000 monoclonal mouse anti-GFP (clones 7.1 and 13.1, 11814460001, Roche), 1:1000 mouse monoclonal anti-V5 (clone V5-10, V8012, Sigma Aldrich), 1:3000 mouse monoclonal anti-GAPDH (sc-47724, Santa Cruz Biotechnology), 1:500 mouse polyclonal anti-M6P (ABCD AG949, ABCD Antibodies), 1:1000 rabbit polyclonal anti-progranulin (18410-1-AP, Proteintech).

### Immunofluorescence

For LysoTracker staining, cells were seeded onto poly-L-lysine-coated glass coverslips and cultured for 24 h. Cells were washed twice with PBS and incubated with LysoTracker (1 μM; Thermo Fisher Scientific) and Hoechst 33342 (1 μg/ml; Bio-Rad) for 20 min at 37°C. Following staining, cells were washed three times with PBS, fixed with 4% paraformaldehyde in PBS for 10 min at room temperature, and mounted in Fluoromount-G mounting medium (Thermo Fisher Scientific). Images were acquired using a Leica SP5 confocal microscope. Image processing and quantification of LysoTracker-positive puncta were performed using Fiji (Schindelin *et al*, 2012).

### RUSH assay

The RUSH assay was performed essentially as previously described (Boncompain *et al*., 2012; Nuchel *et al*., 2026), using a PSAP-RUSH construct as cargo. WI-26 cells were seeded in 12-well plates and transfected with 2 µg of PSAP-RUSH plasmid DNA using X-treme Gene HP transfection reagent (Roche) in a 2:1 DNA/transfection reagent ratio according to the manufacturer’s instructions. Twenty-four hours after transfection, cells were washed once with PBS and incubated in Opti-MEM (Thermo Fisher Scientific) supplemented with 40 µM biotin to synchronously release PSAP from the endoplasmic reticulum. Cells and conditioned media were collected after 0, 2, 4, and 8 h for analysis by western blotting.

### Western blotting

Whole-cell lysates were prepared in SDS sample buffer (50 mM Tris, pH 6.8, 10 mM EDTA, pH 8.0, 5% glycerol, 2% SDS, and 0.01% bromophenol blue) supplemented with 5% 2-mercaptoethanol and incubated at 65°C for 10 min before separation by Tris-glycine SDS-polyacrylamide gel electrophoresis. For PNGase F digestion, lysates were diluted 1:4 with water and incubated with PNGase F (New England Biolabs) or left untreated according to the manufacturer’s instructions prior to electrophoresis. To analyze secreted proteins, cells were cultured in Opti-MEM (Thermo Fisher Scientific) for 24 h before harvesting. Conditioned media were cleared of dead cells and cell debris by sequential centrifugation at 500 x g for 5 min and 16,800 x g for 15 min at 4°C. Proteins were precipitated with 10% trichloroacetic acid for 5 min on ice, collected by centrifugation at 16,800 x g for 15 min at 4°C, washed with acetone, air-dried, and resuspended in SDS sample buffer before electrophoresis. Where indicated, 2,2,2-trichloroethanol (TCE; T54801, Sigma-Aldrich) was incorporated into SDS-polyacrylamide gels. TCE fluorescence was recorded using a ChemiDoc Touch Imaging System (Bio-Rad Laboratories) and served as a loading control. Proteins were transferred onto PVDF membranes by semi-dry blotting. SDS-PAGEs for the RUSH assays were instead blotted by wet blotting onto PVDF membranes. Where indicated, membranes were stained with Ponceau S solution for 5 min after transfer, destained with water, and used as an additional loading control. Following incubation with the indicated primary and secondary antibodies, signals were detected by enhanced chemiluminescence using an ImageQuant 800 Fluor system (Cytiva) controlled by ImageQuant 800 software (version 2.0.0). Signal intensities were quantified using Fiji (Schindelin *et al*., 2012).

### Real-time quantitative reverse transcription and polymerase chain reaction

RNA was isolated using the NucleoSpin RNA kit (Macherey-Nagel) according to the manufacturer’s instructions. cDNA was synthesized using the RevertAid First Strand cDNA synthesis kit (Thermo Fisher Scientific) using random hexamer primers according to the manufacturer’s instructions. The quantitative reverse transcription PCR (qRT-PCR) was set up using SensiFAST SYBR No-ROX kit (Bioline) in technical triplicates. qPCRs were run on the Roche Light Cycler 480 II using the Light Cycler 480 SW software (version 1.51). Primers amplifying human haptoglobin (fwd: 5′-AAGCAGTATGTGGGAAGCCC-3′, rvs: 5′-CCTTTGGCATCCAGGTGTCC-3′), human AXIN2 (fwd: 5′-ACAGGTCGCAGGATGTCTGG-3′, rvs: 5’-TTGTGCTTTGGGCACTATGGG-3′), human WNT3a (fwd: 5-′CCGTGCTGGACA AAGCTACC-3′, rvs: 5-′TCACTGCAAAGGCCACACC-3′), human PSAP (fwd: 5′-CAGAGCTGGACATGACTGAGG-3′, rvs: 5′-CGTCCCCATTATCCTTTGGC-3′), human CatD (fwd: 5′-CCCGCGATCACACTGAAGC-3′, rvs: 5′-GCCTGCGACACCTTGAGC-3′), human PGRN (fwd: 5′-GGTGCCCTGATGGTTCTACC-3′, rvs: 5′-AGCAGGTGGCGTTGGG-3′) and human CatL (fwd: 5′-CCGGGTGGACACAGGTTTTA-3′, rvs: 5′-TCAAATGTTAGAGTAGCTGAGGCA-3′) were used. Relative changes in gene expression were calculated using the 2^ΔΔCt method using the geometric mean of the expression of beta-2-microglobulin (fwd: 5′-CACGTCATCCAGCAGAGAAT-3′, rvs: 5′-TGCTGCTTACATGTCTCGAT-3′), and TATA-binding protein (fwd: 5′-CCGGCTGTTTAACTTCGCTT-3′, rvs: 5′-ACGCCAAGAAACAGTGATGC-3′) for normalization.

### Quantification, structural and statistical analyses

Statistical analyses were performed using GraphPad Prism (version 10.2.0). Quantification of western blot and qRT-PCR data was performed using Microsoft Excel (version 16.95). Structural models were retrieved from the AlphaFold Protein Structure Database (Varadi *et al*, 2022) and visualized using UCSF ChimeraX version 1.9 (Pettersen *et al*, 2021).

## Supporting information

Expanded View Material

## Acknowledgments

We thank Marion Martens and Noah Grohs for technical assistance, Friederike Korn for helpful comments on the manuscript, Thomas Ruppert (ZMBH MS facility), Jan-Wilm Lackmann (CECAD MS facility) for the MS analysis. The work was funded by the Deutsche Forschungsgemeinschaft (DFG, German Research Foundation) - Project-ID 201348542 - CRC 1036/TP12 and the Center of Molecular Medicine Cologne (CMMC). We acknowledge support by Köln Fortune (project 28/2026) to S.S.S.

## Author contributions

SSS: Conceptualization, Investigation, Methodology, Formal analysis, Visualization, Writing – original draft, Writing – review & editing. CN: Investigation, Formal analysis. MT: Investigation, Formal analysis. MP: Investigation. IK: Investigation. JDK: Investigation, Methodology, Formal analysis, Writing – review & editing. JN: Conceptualization, Supervision, Formal analysis, Writing – review & editing. MKL: Conceptualization, Supervision, Funding acquisition, Project administration, Writing – original draft, Writing – review & editing.

