## Supplementary material for "Proteolytic Remodeling of Cargo Receptor Networks by RHBDL4 Tunes Secretory Pathway Flux": Expanded View Material

#### Expanded View Figure Legends

**Figure EV1. RHBDL4 cleaves members of the ERGIC cargo receptor family.**

**(A)** Western blot (WB) analysis of HEK293T cells co-expressing FLAG-ERGIC1 (closed arrow) with either empty vector (-), wild-type (WT) HA-RHBDL4 (HA-R4), or its catalytically inactive SA mutant. WT HA-R4 generates N-terminal cleavage fragments of approximately 22 and 18 kDa (open arrow and open circle), which are stabilized by MG132 (2 µM; DMSO, vehicle control). Overexpression of both HA-R4 constructs also stabilizes the de-glycosylated full-length species (closed circle), phenocopying MG132 treatment and suggesting stabilization of an ERAD dislocation intermediate. Upper panel, schematic of the FLAG-ERGIC1 construct. CHO, N-linked glycosylation site, TMD, TM domain. **(B)** Same experiment as in (A) using FLAG-ERGIC3. WT HA-R4 generates N-terminal FLAG-ERGIC3 cleavage fragment of approximately 50 and 35 kDa (open arrows), which are stabilized by MG132 treatment. Upper panel, schematic of FLAG-ERGIC3 construct. **(C)** WB analysis of FLAG-ERGIC1 (closed arrow) cleavage in HEK293T cells expressing empty vector or WT HA-R4. Inhibition of p97 with CB-5083 (1 µM) stabilizes both the glycosylated (open arrow) and de-glycosylated (open circle) RHBDL4 cleavage products. **(D)** Same experiment as in (C) using FLAG-ERGIC2 as substrate. **(E)** Same experiment as in (C) using FLAG-ERGIC3 as substrate. **(F)** Sanger-sequencing of the ERGIC2 locus confirms correct insertion of the FLAG-sequence in endogenously tagged ERGIC2-FLAG HEK293T cells. **(G)** HEK293T cells were transiently transfected with either empty vector, GFP-tagged RHBDL4 (R4-GFP) WT, the catalytically inactive SA mutant or the SA-UIM double mutant (SAUM) were lysed with Triton X-100 and subjected to GFP-specific immunoprecipitation (IP). WB analysis revealed preferential co-purification of endogenous ERGIC3 (closed arrow) and a ladder of ubiquitinated species (open arrow). Cells were treated with CB-5083 (2.5 µM) prior to lysis.

**Figure EV2. RHBDL4 cleavage site maps to the luminal domains of ERGIC cargo receptors.**

**(A)** Western blot (WB) analysis of HEK293T cells co-expressing full-length (FL) FLAG-ERGIC1 (closed arrow and circle) with either empty vector (-) or wild-type HA-RHBDL4 (HA-R4).

FLAG-ERGIC1 truncation mutants terminating after residues 149 (*149) and 156 (*156) were expressed and compared with RHBDL4-generated cleavage fragments (open arrow and circle) to map the cleavage region. Comparison with the reference peptides localizes the cleavage region to around residue 148, which does not coincide with a predicted rhomboid cleavage motif (see panel C). Samples were left untreated or digested with PNGase F. Cells were treated with MG132 (2 μM) to stabilize cleavage fragments and reference peptides. **(B)** Sample analysis as in (A) using FLAG-ERGIC3 reference peptides. Comparison with the reference peptides maps the cleavage regions to residues 118 and 310, of which only residue 310 is adjacent to a predicted rhomboid cleavage motif (see panel E). **(C)** Schematic of ERGIC1 showing the mapped RHBDL4 cleavage region (indicated by red line) and predicted rhomboid cleavage motifs (Strisovsky *et al.*, 2009). Lower panel, AlphaFold ribbon model of ERGIC1 with the cleavage region highlighted in red and TM domains are shown in grey. **(D)** Schematic of ERGIC2 showing the mapped RHBDL4 cleavage regions (red lines) and predicted rhomboid cleavage motifs. Cleavage regions overlapping predicted rhomboid motifs are indicated by blue arrows. Lower panel, AlphaFold model of ERGIC2 shown as a ribbon representation. The mapped cleavage region is highlighted in red; TM domains are shown in grey. **(E)** Schematic of ERGIC3 showing the mapped RHBDL4 cleavage regions (red lines) and predicted rhomboid cleavage motifs. Cleavage regions overlapping predicted rhomboid motifs by striped arrows. Lower panel, AlphaFold ribbon model of ERGIC3 with cleavage regions highlighted in red and TM domains shown in grey. **(F)** WB analysis of HEK293T cells expressing either wild-type (WT) FLAG-ERGIC2 or an Ala7 mutant in which the predicted cleavage motif surrounding residue 132 was replaced by alanine residues (closed arrow) together with empty vector (-) or WT HA-R4. Mutation of the predicted cleavage motif enhances RHBDL4-dependent cleavage (open arrows and circle). Cells were treated with MG132 (2 μM). **(G)** WB analysis of HEK293T cells expressing WT FLAG-ERGIC2 or the indicated point mutants (S132F, S132P, S132L) (closed arrow) together with empty vector or WT HA-R4. Substitution at the mutated site enhances RHBDL4-dependent cleavage (open arrows and circle). Cells were treated with MG132 (2 μM).

**Figure EV3. RHBDL4 selectively recognizes cargo receptor substrates.**

**(A)** Western blot (WB) analysis of HEK293T cells expressing FLAG-LMAN1 (closed arrow) together with either an empty vector (-), wild-type (WT) HA-RHBDL4 (HA-R4), or the catalytic SA mutant. WT HA-R4 does not generate detectable LMAN1 cleavage fragments, even in the presence of MG132 (2 μM; DMSO, vehicle control). Upper panel, schematic of the FLAG-LMAN1 construct; SP, signal peptide; TMD, TM domain. **(B)** Same experiment as in (A) using the FLAG-LMAN2 construct. Upper panel, schematic of the FLAG-LMAN2 construct; CHO, N-linked glycosylation site. **(C)** Schematic of the TM domain (TMD) charge mutants generated for FLAG-ERGIC2 and FLAG-LMAN2. N, amino terminus, C, carboxyl terminus. **(D)** WB analysis of HEK293T cells expressing either WT FLAG-LMAN2 or the FLAG-LMAN2-I334K mutant (closed arrow) together with empty vector, WT HA-R4, or the catalytic SA mutant. Introduction of a positive charge into the TM domain renders FLAG-LMAN2 susceptible to RHBDL4 cleavage (open arrow). Cells were treated with either DMSO or MG132 (2 μM). **(E)** WB analysis of HEK293T cells co-expressing either WT FLAG-ERGIC2 or the H333L mutant (closed arrow) with empty vector, WT HA-R4, or the catalytic SA mutant. Intensity of all cleavage fragments is reduced for FLAG-ERGIC2-H333L. Cells were treated with MG132 (2 μM) (means $\pm$ SEM, n = 3, ** $p < 0.01$, unpaired two-sided Student's t-test).

**Figure EV4. Functional validation of RHBDL4-dependent ERGIC cargo receptor regulation.**

**(A)** Western blot (WB) analysis of HEK293T cells expressing FLAG-Evi (closed arrow) with either empty vector (-), wild-type (WT) HA-RHBDL4 (HA-R4), or its catalytic SA mutant. WT HA-R4 generates N-terminal FLAG-Evi cleavage fragments (open arrow). Cells were treated with DMSO or MG132 (2 μM). **(B)** Quantification of FLAG-Evi and FLAG-ERGIC2 cleavage based on the ratio of full-length proteins to cleavage fragment intensity (means $\pm$ SEM, n = 3, ** $p < 0.01$, unpaired two-sided Student's t-test). **(C)** WB validation of RHBDL4 knockout (ΔR4) in HCT116 cell clone 2E8. **(D)** TIDE analysis of the RHBDL4 locus in HCT116 ΔR4 clone 2E8 confirms genome editing with an estimated knockout efficiency of 99.3%. **(E)** qRT-PCR analysis showing unchanged *WNT3A* mRNA levels in HCT116 ΔR4 cells (means $\pm$ SEM, n = 5, unpaired two-sided Student's t-test). **(F)** qRT-PCR analysis showing unchanged haptoglobin (*HPT*) mRNA levels following RHBDL4 knockdown in HepG2 cells (means $\pm$ SEM, n = 2).

**Figure EV5. Secretion of lysosomal precursors is increased in RHBDL4 knockout cells.**

**(A)** TIDE analysis of the RHBDL4 locus in HEK293T ΔR4 clone 2B6 (clone 1) and 2G9 (clone 2) confirms knockout efficiency of 90.7% and 92.3%, respectively. **(B)** Western blot (WB)-analysis of endogenous PSAP in wild-type (WT) and RHBDL4 knockout (ΔR4) HEK293T cells. PSAP is detected as two N-glycosylated species (closed arrows), both of which were quantified. Secreted PSAP levels are significantly increased in all three ΔR4 clones, whereas mature saposins are not detected. TCE fluorescence serves as the loading control (means $\pm$ SEM, n = 4, * $p < 0.05$, *** $p < 0.001$, unpaired two-sided Student's t-test). **(C)** Same assay as in (B) analyzing endogenous CatL. Secretion of the CatL pro-form (closed arrow) is increased, whereas intracellular pro-CatL is not detected. The intracellular mature single-chain CatL form is indicated by the grey arrow. Asterisk denotes nonspecific background bands (means $\pm$ SEM, n = 6, * $p < 0.05$, ** $p < 0.01$, unpaired two-sided Student's t-test). **(D)** Same assay as in (B) analyzing endogenous PGRN. Secretion of PGRN (closed arrow) is increased in ΔR4 cells (means ± SEM, n = 4, ** $p < 0.01$, *** $p < 0.001$, unpaired two-sided Student's t-test) **(E)** qRT-PCR analysis of *CATD* mRNA levels in WT and ΔR4 HEK293T cells shows no significant change (means ± SEM, n = 4, unpaired two-sided Student's t-test). **(F)** Sample analysis as in (E) for *PSAP* mRNA levels (means ± SEM, n = 4, unpaired two-sided Student's t-test). **(G)** Sample analysis as in (E) for *CATL* mRNA levels (means ± SEM, n = 4, unpaired two-sided Student's t-test). **(H)** Sample analysis as in (E) for *PGRN* mRNA levels (means ± SEM, n = 4, * $p < 0.05$, unpaired two-sided Student's t-test). **(I)** TIDE analysis of the RHBDL4 locus in ΔR4 WI-26 cells. **(J)** qRT-PCR analysis of CATD and PSAP mRNA levels in WT and ΔR4 WI-26 cells show no significant change (means ± SEM, n = 3, unpaired two-sided Student's t-test).

**Figure EV6. Lysosomal trafficking is preserved in RHBDL4 knockout cells, whereas lysosomal precursor proteins are cleaved by RHBDL4 with different efficiencies.**

**(A)** Quantification of mature CatD heavy-chain levels determined by Western blot (WB) analysis. Mature CatD levels are significantly changed in one RHBDL4 knockout (ΔR4) HEK293T clone (n = 7, means ± SEM, ** $p < 0.01$, unpaired two-sided Student's t-test). **(B)** Quantification of mature single-chain CatL levels determined by WB analysis. Mature CatL levels are significantly changed in one ΔR4 HEK293T clone (n = 5, means ± SEM, ** $p < 0.01$, unpaired two-sided Student's t-test). **(C)** LysoTracker staining of wild-type (WT) and ΔR4 WI-26 cells show no significant difference in the number of LysoTracker-positive puncta per cell. Scale bar equals 10 µm, (n = 10, means ± SEM, unpaired two-sided Student's t-test) **(D)** WB analysis of HEK239T cells expressing CatD-FLAG (closed arrow) together with either empty vector (-), WT HA-RHBDL4 (HA-R4), or the catalytic SA mutant. WT HA-R4 generates a CatD cleavage fragment (open arrow). Cells were treated with either DMSO or MG132 (2 µM). **(E)** Same assay as in (D) using CatL-FLAG. **(F)** Same assay as in (D) using PSAP-FLAG. **(G)** Quantification of PSAP-FLAG, CatD-FLAG, and CatL-FLAG cleavage based on the ratio of full-length proteins to cleavage fragment intensity (means ± SEM). **(H)** WB analysis of M6P-modified proteins (closed arrows) in WT and ΔR4 HEK293T cells. M6P levels are unchanged in ΔR4 cells. Lysates from GNPTAB knockout (ΔGNPTAB) cells serve as specificity control for the M6P signal. TCE fluorescence serves as the loading control.

**Figure EV7. RHBDL4 antagonizes sortilin secretion.**

Western blot (WB) analysis of sortilin secretion in HEK293T cells expressing sortilin together with either empty vector (-) or wild-type (WT) HA-RHBDL4 (HA-R4). Where indicated, cells were treated with ADAM metalloprotease inhibitor BB94 (10 µM). Ectopically expressed sortilin is secreted as both the full-length protein and a shed ectodomain. WT HA-R4 reduces secretion of both forms. Asterisk indicates nonspecific background bands.

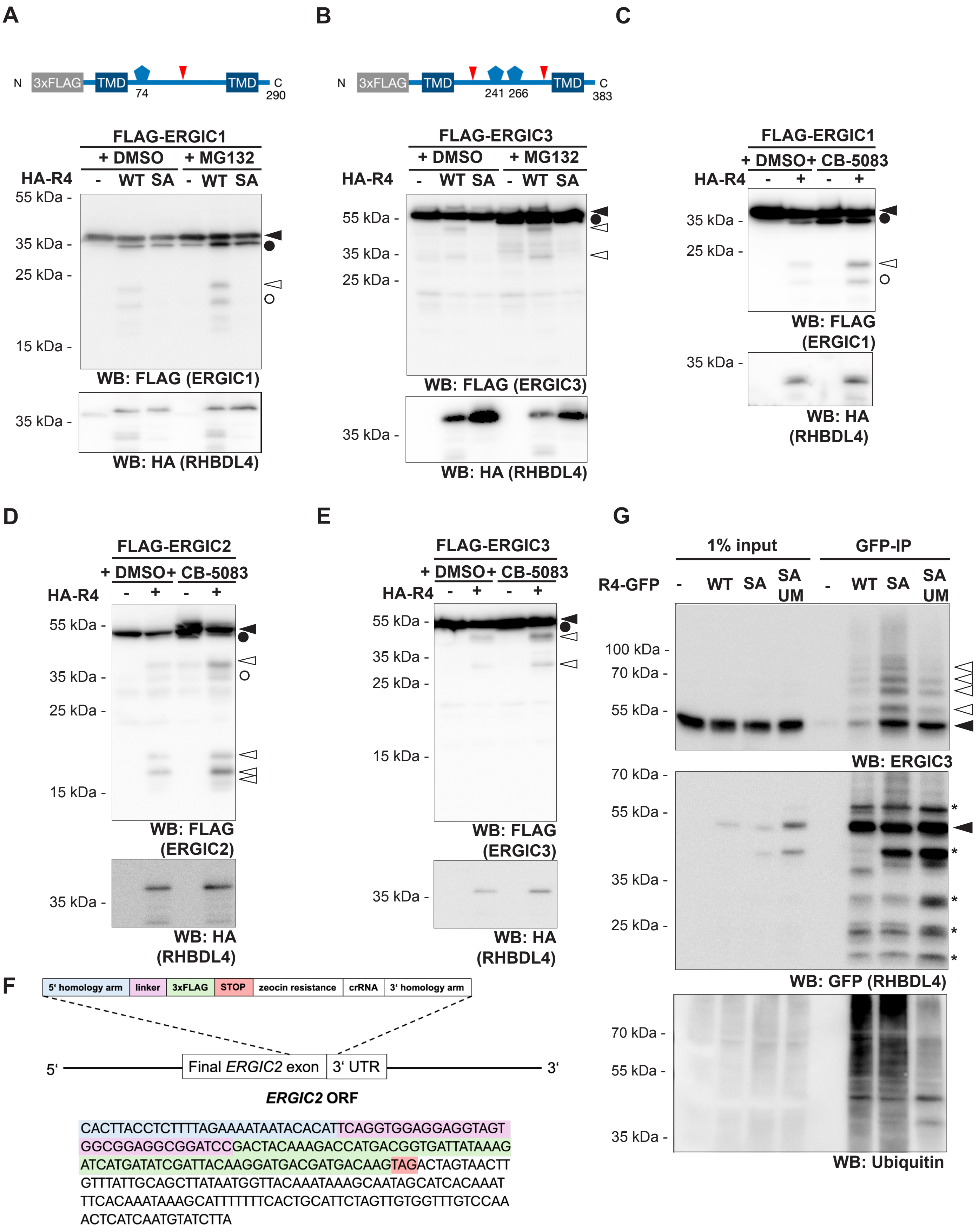
A
N 3xFLAG TMD 74 TMD C 290
FLAG-ERGIC1
+ DMSO + MG132
HA-R4 - WT SA - WT SA
55 kDa -
35 kDa -
25 kDa -
15 kDa -
WB: FLAG (ERGIC1)
35 kDa -
WB: HA (RHBDL4)
B
N 3xFLAG TMD 241 266 TMD C 383
FLAG-ERGIC3
+ DMSO + MG132
HA-R4 - WT SA - WT SA
55 kDa -
35 kDa -
25 kDa -
WB: FLAG (ERGIC3)
35 kDa -
WB: HA (RHBDL4)
C
FLAG-ERGIC1
+ DMSO + CB-5083
HA-R4 - + - +
35 kDa -
25 kDa -
WB: FLAG (ERGIC1)
35 kDa -
WB: HA (RHBDL4)
D
FLAG-ERGIC2
+ DMSO + CB-5083
HA-R4 - + - +
55 kDa -
35 kDa -
25 kDa -
15 kDa -
WB: FLAG (ERGIC2)
35 kDa -
WB: HA (RHBDL4)
E
FLAG-ERGIC3
+ DMSO + CB-5083
HA-R4 - + - +
55 kDa -
35 kDa -
25 kDa -
15 kDa -
WB: FLAG (ERGIC3)
35 kDa -
WB: HA (RHBDL4)
G
1% input
GFP-IP
R4-GFP - WT SA SA UM - WT SA SA UM
100 kDa -
70 kDa -
55 kDa -
WB: ERGIC3
70 kDa -
55 kDa -
35 kDa -
25 kDa -
*
WB: GFP (RHBDL4)
70 kDa -
55 kDa -
35 kDa -
WB: Ubiquitin
F
5‘ homology arm
linker
3xFLAG
STOP
zeocin resistance
crRNA
3‘ homology arm
5‘
Final ERGIC2 exon
3‘ UTR
3‘
ERGIC2 ORF
CACTTACCTCTTTTAGAAAATAATACACATTCAGGTGGAGGAGGTAGT
GGCGGAGGCGGATCCGACTACAAAGACCATGACGGTGATTATAAAG
ATCATGATATCGATTACAAGGATGACGATGACAAGTAGACTAGTAACTT
GTTTATTGCAGCTTATAATGGTTACAAATAAAGCAATAGCATCACAAAT
TTCACAAATAAAGCATTTTTTTCACTGCATTCTAGTTGTGGTTTGTCCAA
ACTCATCAATGTATCTTA

**Steigleder *et al.*, EV2**

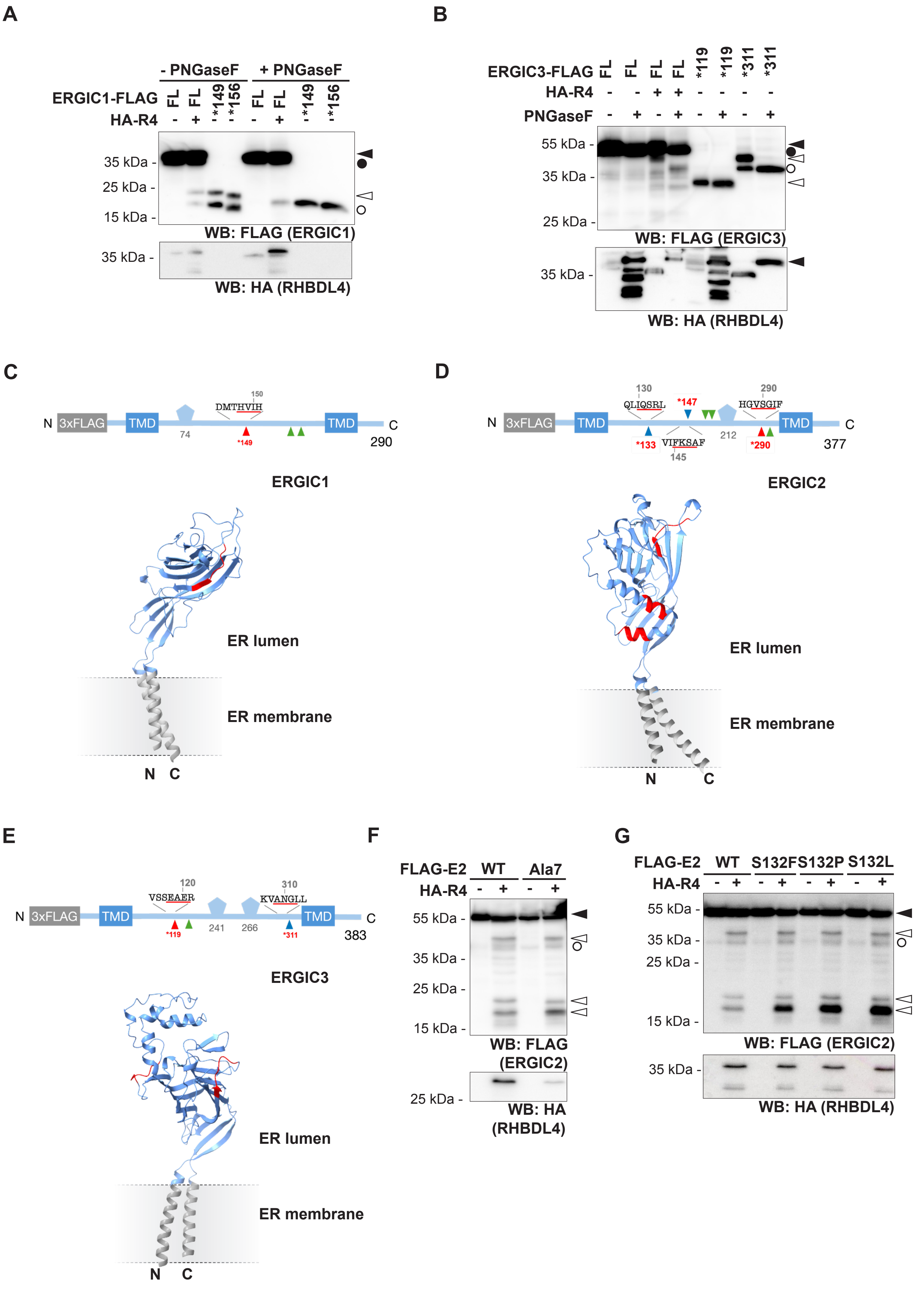

Steigleder *et al.*, EV3

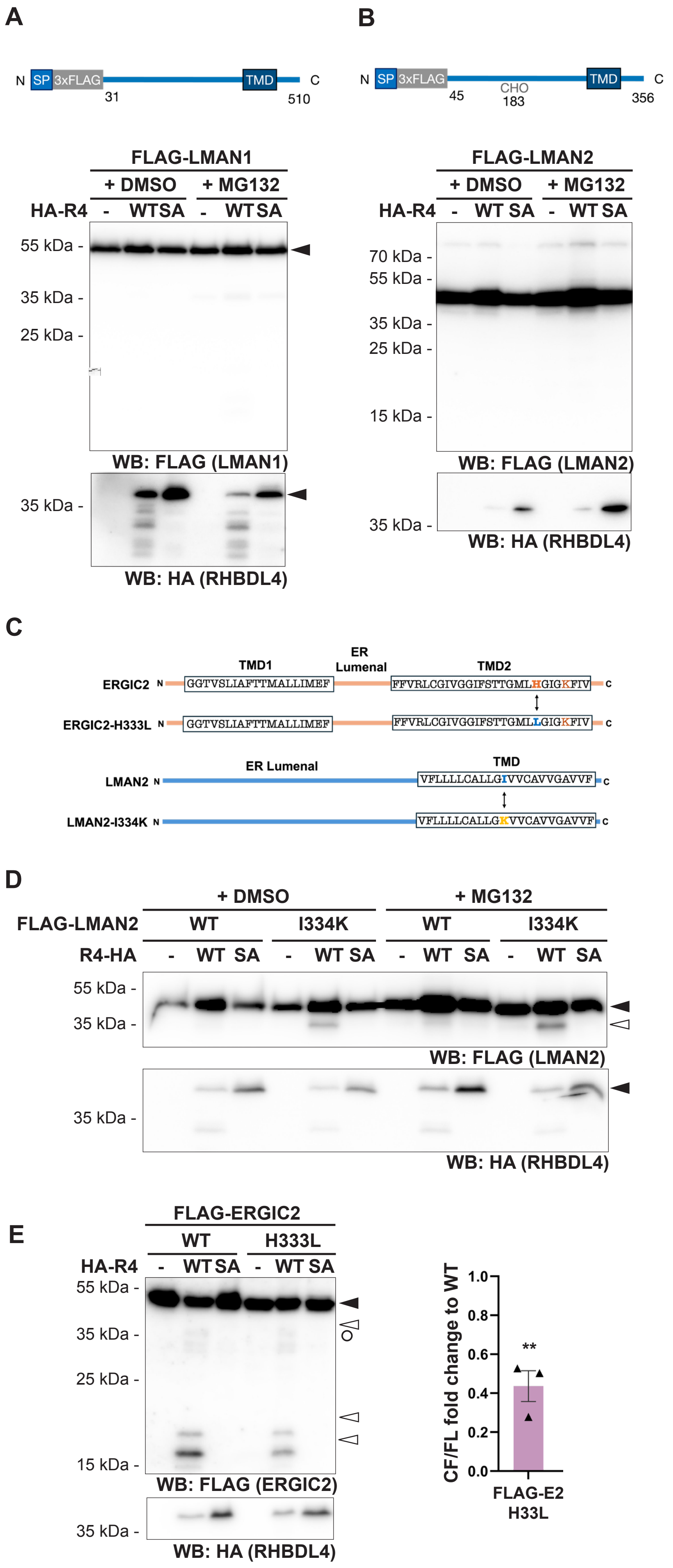

**Steigleder *et al.*, EV4**

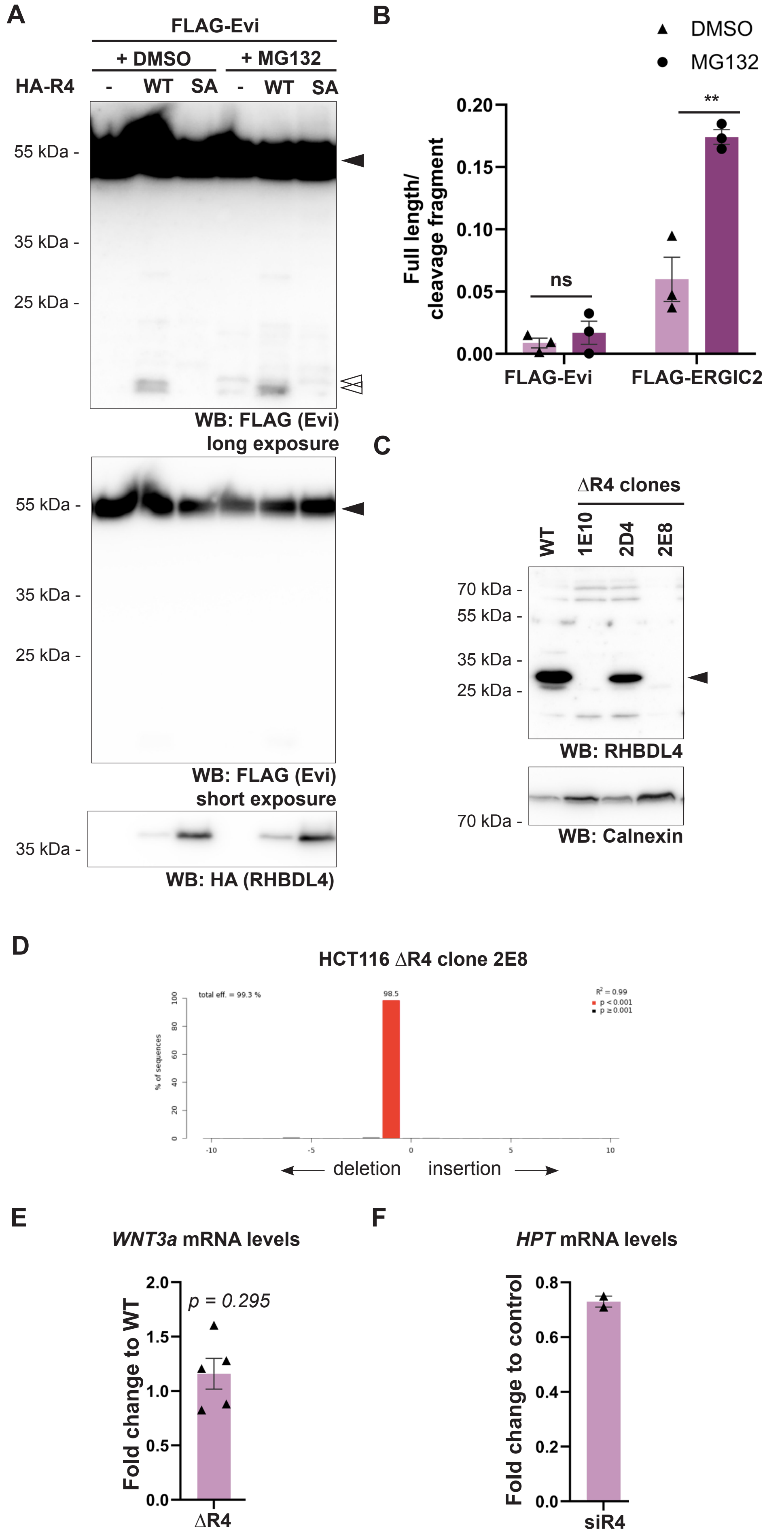

Steigleder *et al.*, EV5

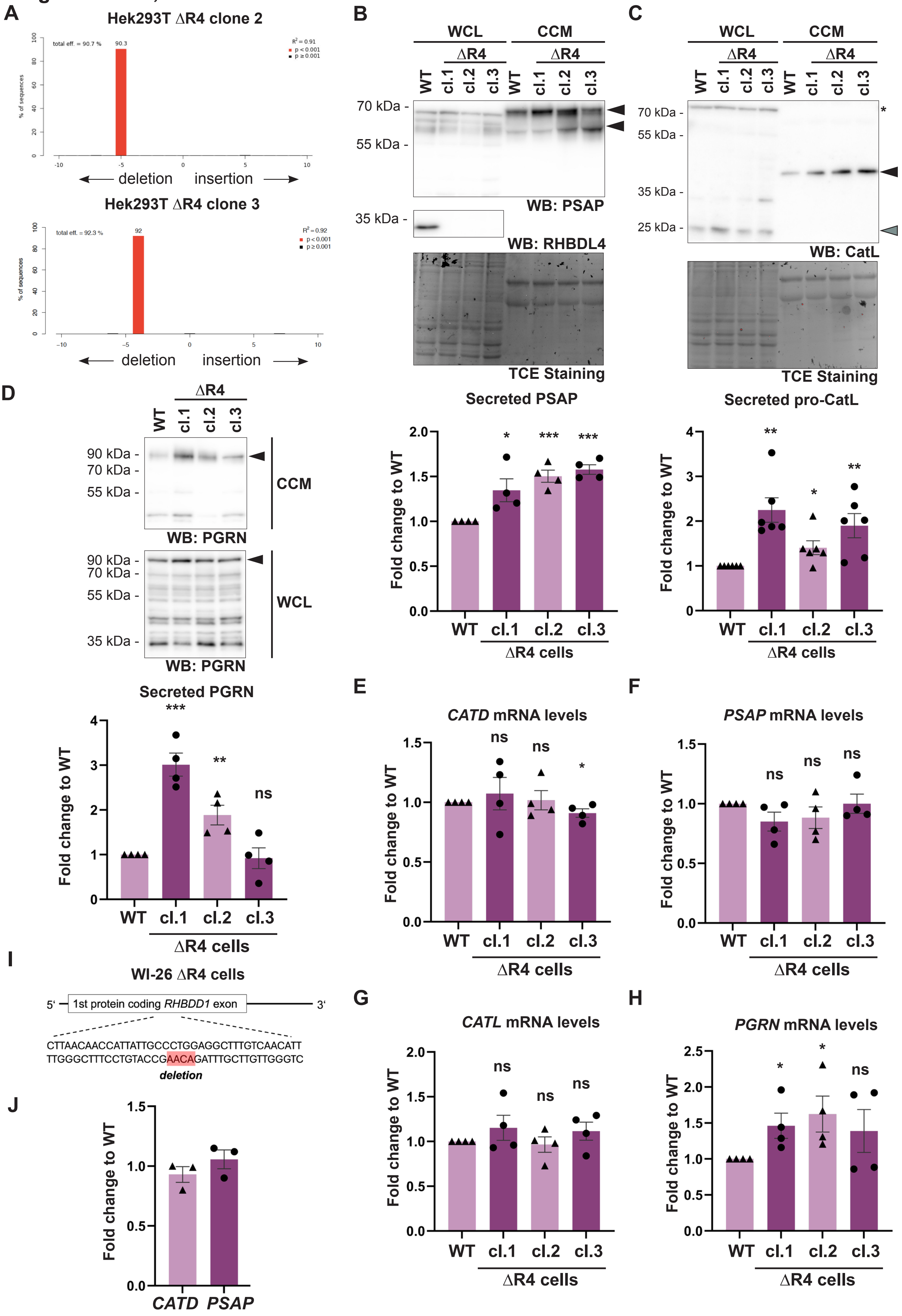

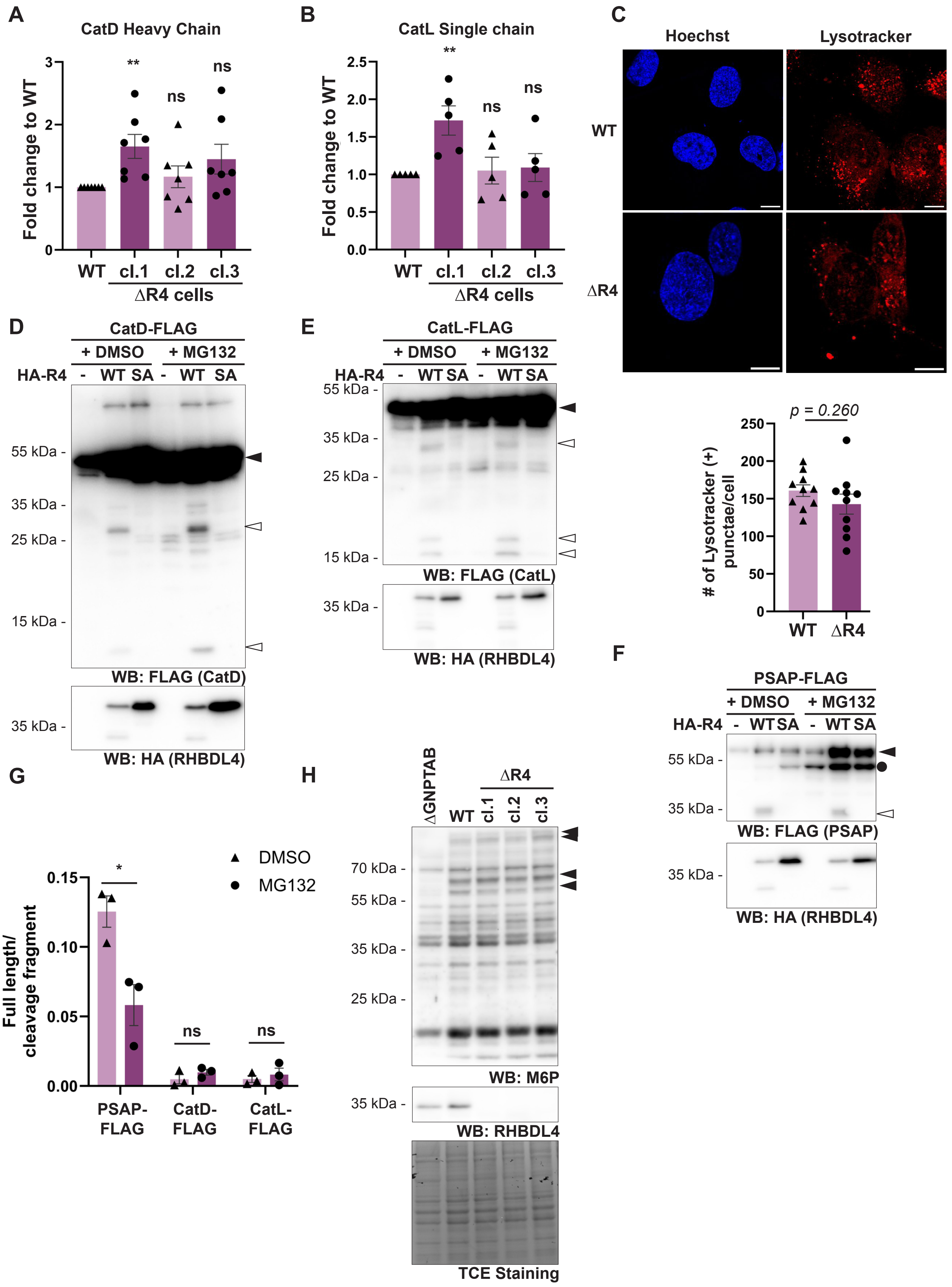
A
CatD Heavy Chain
Fold change to WT
**
ns
ns
WT cl.1 cl.2 cl.3
ΔR4 cells
B
CatL Single chain
**
ns
ns
WT cl.1 cl.2 cl.3
ΔR4 cells
C
Hoechst
Lysotracker
WT
ΔR4
p = 0.260
### of Lysotracker (+) punctae/cell
WT ΔR4
D
CatD-FLAG
+ DMSO + MG132
HA-R4 - WT SA - WT SA
55 kDa -
35 kDa -
25 kDa -
15 kDa -
WB: FLAG (CatD)
35 kDa -
WB: HA (RHBDL4)
E
CatL-FLAG
+ DMSO + MG132
HA-R4 - WT SA - WT SA
55 kDa -
35 kDa -
25 kDa -
15 kDa -
WB: FLAG (CatL)
35 kDa -
WB: HA (RHBDL4)
F
PSAP-FLAG
+ DMSO + MG132
HA-R4 - WT SA - WT SA
55 kDa -
35 kDa -
WB: FLAG (PSAP)
35 kDa -
WB: HA (RHBDL4)
G
DMSO
MG132
Full length/ cleavage fragment
*
ns
ns
PSAP-FLAG
CatD-FLAG
CatL-FLAG
H
ΔGNPTAB
WT
ΔR4
cl.1 cl.2 cl.3
70 kDa -
55 kDa -
35 kDa -
25 kDa -
WB: M6P
35 kDa -
WB: RHBDL4
TCE Staining

**Steigleder *et al.*, EV7**

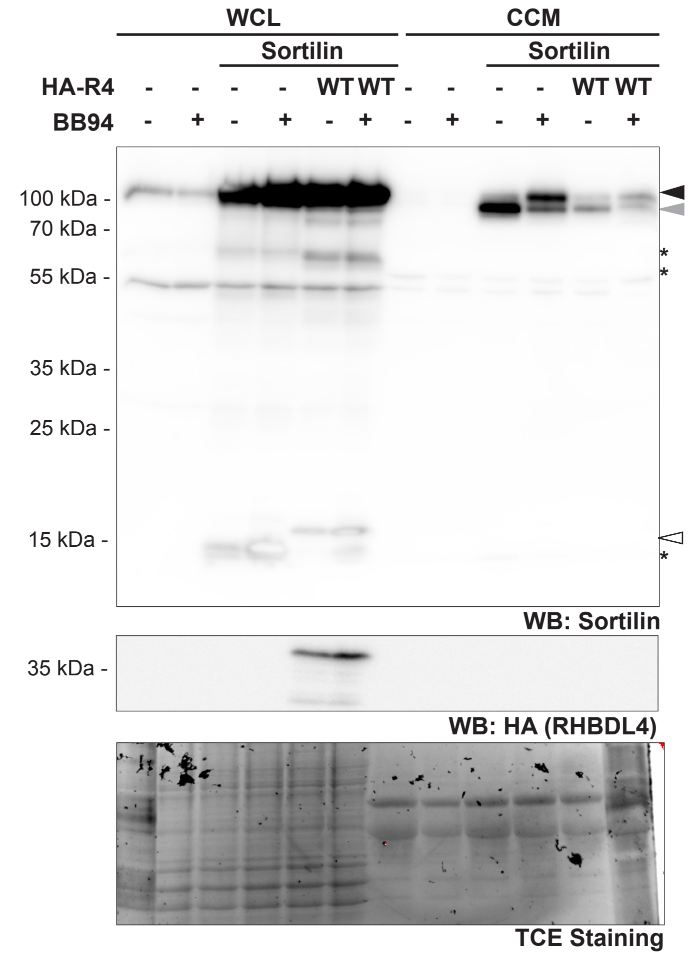

**Table EV1.** Results of the SILAC-based quantitative proteomic screen for endogenous RHBDL4 substrate comparing wild-type (medium) and RHBDL4 knockout (heavy) cells.

| # | Gene Name | Ratio (Heavy/Medium) n = 2 |
|---|---|---|
| 1 | SLC33A1 | 2.07 |
| 2 | SPCS3 | 1.56 |
| 3 | ALG2 | 1.40 |
| 4 | EGFR | 1.39 |
| 5 | DNAJC1 | 1.34 |
| 6 | DOLPP1 | 1.34 |
| 7 | SERINC1 | 1.31 |
| 8 | **ERGIC1** | **1.29** |
| 9 | HLA-B | 1.28 |
| 10 | TMEM109 | 1.26 |
| 11 | MPDU1 | 1.25 |
| 12 | GOLGB1 | 1.23 |
| 13 | LRRC59 | 1.23 |
| 14 | MAN1B1 | 1.20 |
| 15 | B3GALTL | 1.20 |
| 16 | CKAP4 | 1.20 |
| 17 | SRPRB | 1.18 |
| 18 | ALG11 | 1.17 |
| 19 | PNPLA8 | 1.17 |
| 20 | MSMO1 | 1.16 |
| 21 | LPCAT2 | 1.16 |
| 22 | IKBIP | 1.15 |
| 23 | AGPAT1 | 1.14 |
| 24 | LCLAT1 | 1.14 |
| 25 | FADS1 | 1.14 |
| 26 | **ERGIC3** | **1.14** |
| 27 | CDS2 | 1.13 |
| 28 | CANX | 1.13 |
| 29 | HLA-C | 1.13 |
| 30 | TMED9 | 1.12 |
| 31 | TMEM50A | 1.12 |
| 32 | CERS5 | 1.11 |
| 33 | TMED4 | 1.11 |
| 34 | DAD1 | 1.11 |
| 35 | HLA-1 | 1.10 |
| 36 | TMED7 | 1.10 |
| 37 | KTN1 | 1.10 |
| 38 | ALG3 | 1.10 |

**Table EV2:** Proteins identified in all four biological replicates of the RHBDL4 knockout secretome analysis that were enriched by at least 20% compared with wild-type cells.

| **Proteins identified** | **Ratio (Heavy/Medium)** | **p Value** |
|---|---|---|
| COL11A1 | 7.81 | 0.01659 |
| SCG3 | 4.09 | 0.00012 |
| CHGB | 3.33 | 0.00394 |
| **GRN** | **2.94** | **0.00011** |
| GDF7 | 2.6 | 0.00066 |
| CHAD | 2.57 | 0.18122 |
| LYG2 | 2.34 | 0.00054 |
| **CTSD** | **2.19** | **0.00006** |
| ATP6AP2 | 2.15 | 0.00023 |
| BMP7 | 2.08 | 0.00003 |
| FBN1 | 2.00 | 0.00008 |
| MMP2 | 2.00 | 0.00784 |
| LMAN2 | 1.90 | 0.00024 |
| IGFBP4 | 1.82 | 0.00014 |
| PXDN | 1.81 | 0.00008 |
| FABP5 | 1.76 | 0.00293 |
| RNASE4 | 1.76 | 0.00037 |
| MGAT5 | 1.75 | 0.00071 |
| **GM2A** | **1.74** | **0.00004** |
| GLO1 | 1.73 | 0.00188 |
| **PSAP** | **1.67** | **0.0006** |
| NELL2 | 1.65 | 0.00048 |
| **CTSL** | **1.60** | **0.00039** |
| DSP | 1.58 | 0.08925 |
| CHST14 | 1.54 | 0.00413 |
| TLN1 | 1.52 | 0.54542 |
| PCOLCE2 | 1.49 | 0.00029 |
| SPON1 | 1.49 | 0.00222 |
| **LGMN** | **1.48** | **0.00005** |
| HSPG2 | 1.45 | 0.01301 |
| PRNP | 1.44 | 0.00027 |
| LTBP3 | 1.44 | 0.00102 |
| IGF2R | 1.41 | 0.03001 |
| VCL | 1.38 | 0.00897 |
| GAPDH | 1.36 | 0.01521 |
| TIMP2 | 1.34 | 0.00042 |
| GOLM1 | 1.33 | 0.02436 |
| APLP2 | 1.32 | 0.04459 |
| PFN1 | 1.31 | 0.0061 |
| CLEC11A | 1.30 | 0.15452 |
| PPIA | 1.29 | 0.03254 |
| CILP2 | 1.29 | 0.00214 |
| HSPA13 | 1.29 | 0.01551 |
| NUCB1 | 1.28 | 0.00659 |
| NUCB2 | 1.28 | 0.00328 |
| AHCY | 1.25 | 0.64536 |
| LDHB | 1.25 | 0.53341 |
| PTPRF | 1.24 | 0.01642 |
| TPI1 | 1.24 | 0.02974 |

| | | |
|---|---|---|
| HSPA1B;HSPA1A | 1.22 | 0.00247 |
| FSTL1 | 1.22 | 0.00227 |
| NME1 | 1.21 | 0.03436 |
| HTRA1 | 1.21 | 0.00165 |
| TXN | 1.20 | 0.08823 |
